# Mathematical constraints on *F_ST_*: biallelic markers in arbitrarily many populations

**DOI:** 10.1101/094433

**Authors:** Nicolas Alcala, Noah A Rosenberg

**Affiliations:** Department of Biology, Stanford University, Stanford, CA 94305-5020, USA

**Author notes:** Correspondence to Nicolas Alcala.

**Keywords:** allele frequency, *F_ST_*, genetic differentiation, migration, population structure

## Abstract

*F_ST_* is one of the most widely used statistics in population genetics. Recent mathematical studies have identified constraints on *F_ST_* that challenge interpretations of *F_ST_* as a measure with potential to range from 0 for genetically similar populations to 1 for divergent populations. We generalize results obtained for population pairs to arbitrarily many populations, characterizing the mathematical relationship between *F_ST_*, the frequency *M* of the more frequent allele at a polymorphic biallelic marker, and the number of subpopulations *K*. We show that for fixed *K*, *F_ST_* has a peculiar constraint as a function of *M*, with a maximum of 1 only if *M* = *i*/*K* for integers *i* with ⌈*K*/2⌉ ≤ *i* ≤ *K* − 1. For fixed *M*, as *K* grows large, the range of *F_ST_* becomes the full closed or half-open unit interval. For fixed *K*, however, some *M* < (*K* − 1)/*K* always exists at which the upper bound on *F_ST_* is constrained to be below 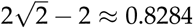. In each of three migration models—island, rectangular stepping-stone, and linear stepping-stone—we use coalescent simulations to show that under weak migration, *F_ST_* depends strongly on the allele frequency *M* when *K* is small, but not when *K* is large. Finally, using data on human genetic variation, we employ our results to explain the generally smaller *F_ST_* values between pairs of continents relative to global *F_ST_* values. We discuss implications for the interpretation and use of *F_ST_*.

---

GENETIC differentiation, in which individuals from the same subpopulation are more genetically similar than are individuals from different subpopulations, is a central concept in population genetics. It can arise from a large variety of processes, including from aspects of the physical environment such as geographic barriers and variable permeability to migrants, as well as from biotic phenomena such as assortative mating and self-fertilization. Genetic differentiation among populations is thus a pervasive feature of population-genetic variation.

To measure genetic differentiation, Wright (1951) introduced the fixation index *F_ST_*, defined as the *“correlation between random gametes, drawn from the same subpopulation, relative to the total.”* Many definitions of *F_ST_* and related statistics have since been proposed (reviewed by Holsinger and Weir 2009). For a polymorphic biallelic marker, *F_ST_* is often defined as a ratio of among-subpopulation variance 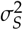 in the frequency of a specific allele A to the “total variance” 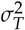 (Weir 1996):

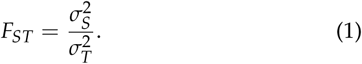

Denoting by *p_k_* the frequency of allele A in subpopulation *k* and by *M* the mean frequency of allele A across all *K* subpopulations, 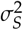 is defined as the variance of *p_k_* across subpopulations, 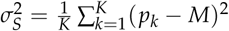. The total variance 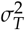 is defined as the variance in an indicator of allelic state for an allele randomly drawn from the entire population, in other words, the variance of a Bernoulli variable with mean *M*, 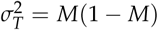. Because by assumption the locus is polymorphic, *M* ≠ 1 and 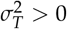.

*F_ST_* and related statistics have a wide range of applications. For example, *F_ST_* is used as a descriptive statistic whose values are routinely reported in empirical population-genetic studies (Holsinger and Weir 2009). It is employed as a test statistic for spatially divergent selection, either acting on a locus (Lewontin and Krakauer 1973; Bonhomme *et al.* 2010) or, using comparisons to a corresponding phenotypic statistic *Q_ST_*, on a trait (Leinonen *et al.* 2013). *F_ST_* is also used as a summary statistic for demographic inference, to measure gene flow between subpopulations (Slatkin 1985), or via approximate Bayesian computation, to estimate demographic parameters (Cornuet *et al.* 2008).

Applications of *F_ST_* generally assume that values near 0 indicate that there are almost no genetic differences among subpopulations, and that values near 1 indicate that subpopulations are genetically different (Hartl and Clark 1997; Frankham *et al.* 2002; Holsinger and Weir 2009). Mathematical studies, however, have challenged the simplicity of this interpretation, commenting that the range of values that *F_ST_* can take is considerably restricted by the allele frequency distribution (Table 1). Such studies have highlighted a direct relationship between allele frequencies and constraints on the range of *F_ST_*, through functions of the allele frequency distribution such as the mean heterozygosity across subpopulations, *H_S_*. The maximal *F_ST_* has been shown to decrease as a function of *H_S_*, both for an infinite (Hedrick 1999) and for a fixed finite number of subpopulations *K* ≥ 2 (Long and Kittles 2003; Hedrick 2005). Consequently, if subpopulations differ in their alleles but separately have high heterozygosity, then *H_S_* can be high and *F_ST_* can be low; *F_ST_* can be near 0 even if subpopulations are completely genetically different in the sense that no allele occurs in more than one subpopulation.

**Table 1.**
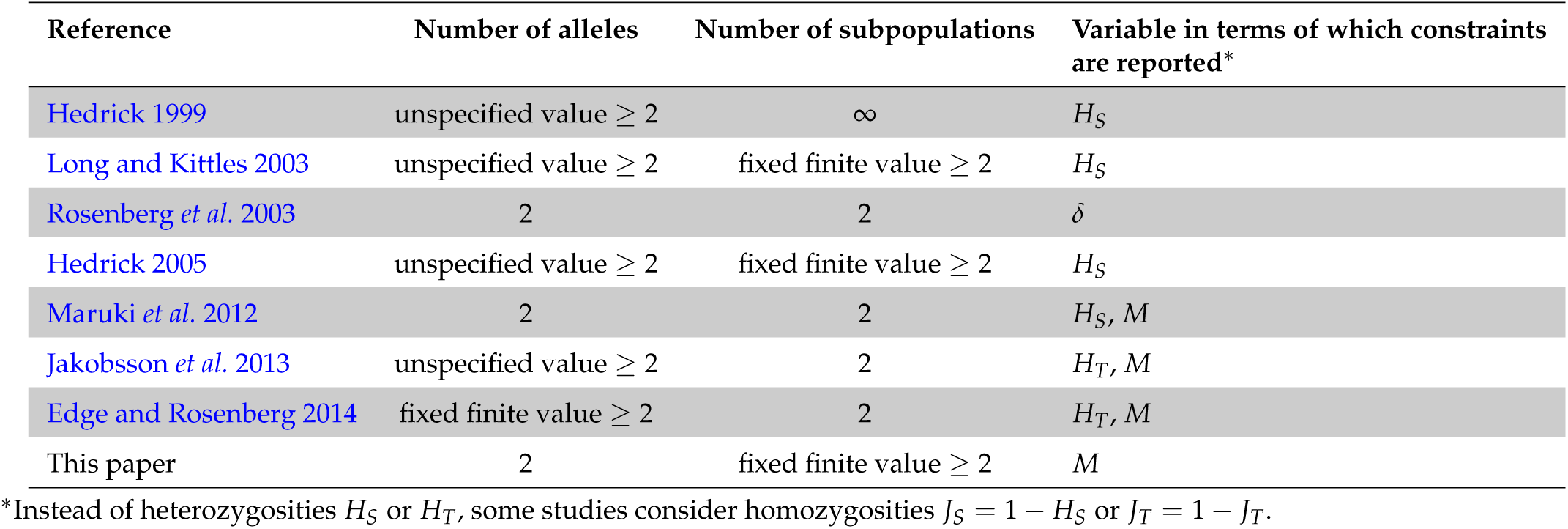
Studies describing the mathematical constraints on *F_ST_*. *H_S_* and *H_T_* denote the within-subpopulation and total heterozygosities, respectively, *δ* denotes the absolute difference in the frequency of a specific allele between two subpopulations, and *M* denotes the frequency of the most frequent allele.

| Reference | Number of alleles | Number of subpopulations | Variable in terms of which constraints are reported* |
| --- | --- | --- | --- |
| Hedrick 1999 | unspecified value $\geq 2$ | ∞ | $H_S$ |
| Long and Kittles 2003 | unspecified value $\geq 2$ | fixed finite value $\geq 2$ | $H_S$ |
| Rosenberg *et al.* 2003 | 2 | 2 | $\delta$ |
| Hedrick 2005 | unspecified value $\geq 2$ | fixed finite value $\geq 2$ | $H_S$ |
| Maruki *et al.* 2012 | 2 | 2 | $H_S, M$ |
| Jakobsson *et al.* 2013 | unspecified value $\geq 2$ | 2 | $H_T, M$ |
| Edge and Rosenberg 2014 | fixed finite value $\geq 2$ | 2 | $H_T, M$ |
| This paper | 2 | fixed finite value $\geq 2$ | $M$ |

\*Instead of heterozygosities $H_S$ or $H_T$ , some studies consider homozygosities $J_S = 1 - H_S$ or $J_T = 1 - H_T$ .

Detailed mathematical results have clarified the relationship between allele frequencies and *F_ST_* in the case of *K* = 2 subpopulations. Considering a biallelic marker, Maruki *et al.* (2012) evaluated the constraint on *F_ST_* by the frequency *M* of the most frequent allele: the maximal *F_ST_* decreases monotonically from 1 to 0 with increasing *M*, 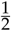 ≤ *M* < 1. Jakobsson *et al.* (2013) extended this result to multiallelic markers with an unspecified number of distinct alleles, showing that the maximal *F_ST_* increases from 0 to 1 as a function of *M* when 0 < *M* < 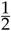, and decreases from 1 to 0 when 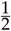 ≤ *M* < 1 in the manner reported by Maruki *et al.* (2012). Edge and Rosenberg (2014) generalized these results to the case of a fixed finite number of alleles, showing that the maximal *F_ST_* differs slightly from the unspecified case when the fixed number of distinct alleles is odd.

In this study, we characterize the relationship between *F_ST_* and the frequency *M* of the most frequent allele, for a biallelic marker and an *arbitrary* number of subpopulations *K*. We derive the mathematical upper bound on *F_ST_* in terms of the frequency *M* of the most frequent allele, extending the biallelic 2-subpopulation result to arbitrary *K*. To assist in interpreting the bound, we simulate the joint distribution of *F_ST_* and *M* in three migration models, describing its properties as a function of the number of subpopulations and the migration rate. The *K*-population upper bound on *F_ST_* as a function of *M* facilitates an explanation of counterintuitive aspects of global human genetic differentiation. We discuss the importance of the results for applications of *F_ST_* more generally.

## Mathematical constraints

### Model

Our goal is to derive the range of values *F_ST_* can take—the lower and upper bounds on *F_ST_*—as a function of the frequency *M* of the most frequent allele for a biallelic marker when the number of subpopulations *K* is a fixed finite value greater than or equal to 2. We consider a polymorphic locus with two alleles, A and a, in a setting with *K* subpopulations contributing equally to the total population. We denote the frequency of allele A in subpopulation *k* by *p_k_*. The frequency of allele a in subpopulation *k* is 1 − *p_k_*. Each allele frequency *p_k_* lies in the interval [0,1].

The mean frequency of allele A across the subpopulations is 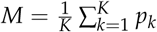, and the mean frequency of allele a is 1 − *M*. Without loss of generality, we assume that allele A is the more frequent allele in the total population, so that *M* ≥ 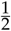 ≥ 1 − *M*. Because by assumption the locus is polymorphic, *M* ≠ 1.

We assume that the allele frequencies *M* and *p_k_* are parametric allele frequencies of the total population and subpopulations, and not estimated values computed from data. For simplicity, we hereafter refer to *F_ST_* as *F*.

### F as a function of M

Eq. 1 expresses *F* as a ratio of among-subpopulation variance, 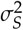, to total variance in allele frequency, 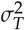. We can write *F* in terms of allele frequencies by substituting 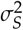 and 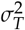 in eq. 1 with their respective expressions in terms of allele frequencies:

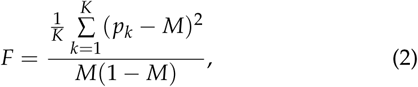

where *p_k_* ranges in [0,1] and *M* ranges in [ 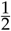, 1).

Note that this expression is equivalent to the expression for *F_ST_* in the *G_ST_* framework of Nei (1973), also used in eq. 1 of Jakobsson *et al.* (2013) and eq. 1 of Edge and Rosenberg (2014). In that context, *G_ST_* is written in terms of a ratio of the mean within-subpopulation heterozygosity, 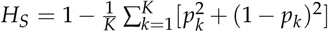, to the total heterozygosity, *H_T_* = 1 − [*M*^2^ + (1 − *M*)^2^]:

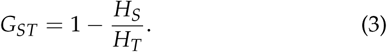

Simplifying eq. 2 by noting that 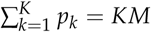 leads to:

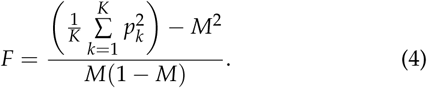

For fixed M, we seek the vectors (*p*_1_, *p*_2_,&, *p_k_*.), with *p_k_* ∈ [0,1] and 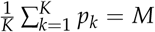, that minimize and maximize *F*.

### Lower bound

From eq. 4, for all *M* ∈ [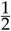,1), setting *p_k_* = *M* in all subpopulations *k* yields *F* = 0. The Cauchy-Schwarz inequality guarantees that 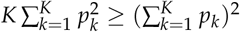, with equality if and only if *p*_1_ = *p*_2_ = … = *p_K_*. Hence 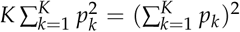, or, dividing both sides by *K*^2^, 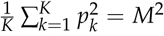, requires *p*_1_ = *p*_2_ = … = *p_k_*. = *M*. Examining eq. 4, (*p*_1_, *p*_2_,…, *p_k_*.) = (*M*, *M*,…, *M*) is thus the only vector that yields *F* = 0. We can conclude that the lower bound on *F* is equal to 0 irrespective of *M*, for any value of the number of subpopulations *K*.

### Upper bound

To derive the upper bound on *F* in terms of *M*, we must maximize *F* in eq. 4, assuming that *M* and *K* are constant. Because all terms in eq. 4 depend only on *M* and *K* except the positive term 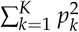 in the numerator, maximizing *F* corresponds to maximizing 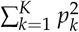 at fixed *M* and *K*.

Denote by [*x*] the greatest integer less than or equal to *x*, and by {*x*} = *x* − [*x*] the fractional part of *x*. Using a result from Rosenberg and Jakobsson (2008), Theorem 1 from Appendix A states that the maximum for 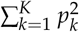 satisfies

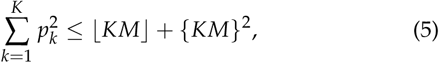

with equality if and only if allele A has frequency 1 in ⌊*KM*⌋ subpopulations, frequency {*KM*} in a single subpopulation, and frequency 0 in all other subpopulations. Substituting eq. 5 into eq. 4, we obtain the upper bound for *F*:

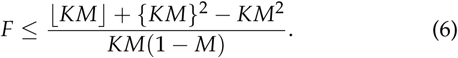

The upper bound on *F* in terms of *M* has a piecewise structure, with changes in shape occurring when *KM* is an integer.

For *i* = ⌊*K*/2⌋, ⌊*K*/2⌋ + 1,…,*K* − 1, define the interval *I_i_*, by 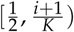 for *i* = [*K*/2] in the case that *K* is odd and by 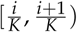 for all other (*i*, *K*). For *M* ∈ *I_i_*, ⌊*KM*⌋ has a constant value *i*. Writing *x* = *KM* − ⌊*KM*⌋ = *KM* − *i* so that 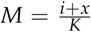, for each interval *I_i_*, the upper bound on *F* is a smooth function

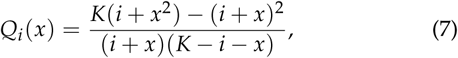

where *x* lies in [0,1) (or in [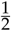, 1) for odd *K* and *i* = ⌊*K*/2⌋), and *i* lies in 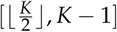.

The conditions under which the upper bound is reached illuminate its interpretation. The maximum requires the most frequent allele to have frequency 1 or 0 in all except possibly one subpopulation, so that the locus is polymorphic in at most a single subpopulation. Thus, *F* is maximal when fixation is achieved in as many subpopulations as possible.

Figure 1 shows the upper bound on *F* in terms of *M* for various values of *K*. It has peaks at values 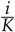, where it is possible for the allele to be fixed in all *K* subpopulations and for *F* to reach a value of 1. Between 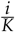 and 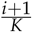, the function reaches a local minimum, eventually decreasing to 0 as *M* approaches 1. The upper bound is not differentiable at the peaks, and it is smooth and strictly below 1 between the peaks. If *K* is even, the upper bound begins from a local maximum at *M* = 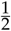, whereas if *K* is odd, it begins from a local minimum at *M* = 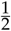.

**Figure 1.**
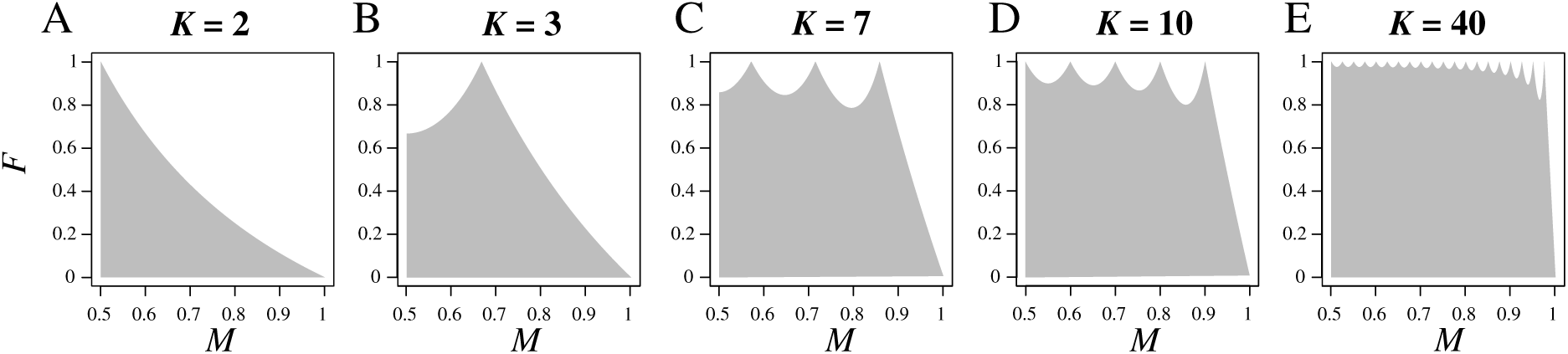
Bounds on *F* as a function of the frequency of the most frequent allele, *M*, for different numbers of subpopulations *K*. (A) *K* = 2. (B) *K* = 3. (C) *K* = 7. (D) *K* = 10. (E) *K* = 40. The shaded region represents the space between the upper and lower bounds on *F*. The upper bound is computed from eq. 6; for each *K*, the lower bound is 0 for all values of *M*.

### Properties of the upper bound

#### Local maxima

We explore properties of the upper bound on *F* as a function of *M* for fixed *K* by examining the local maxima and minima. The upper bound is equal to 1 on interval *I_i_* if and only if the numerator and denominator in eq. 7 are equal. Noting that *K* ≥ 2, this condition is equivalent to *x*^2^ = *x* and hence, because 0 ≤ *x* < 1, *x* = {*KM*} = 0. Thus, on interval *I_i_* for *M*, the maximal *F* is 1 if and only if *KM* is an integer.

*KM* has exactly 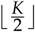 integer values for *M* ∈ [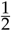,1). Consequently, given *K*, there are exactly 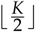 maxima at which *F* can equal 1, at 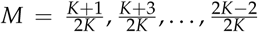 if *K* is odd and at 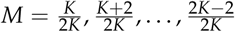 if *K* is even.

This analysis finds that *F* is only unconstrained within the unit interval for a finite set of values of the frequency *M* of the most frequent allele. The size of this set increases with the number of subpopulations *K*.

#### Local minima

Equality of the upper bound at the right endpoint of each interval *I_i_* and the left endpoint of *I*_*i*+1_ for each *i* from 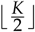 to *K* − 2 demonstrates that the upper bound on *F* is a continuous function of *M*. Consequently, local minima necessarily occur between the local maxima. If *K* is even, then the upper bound on *F* possesses 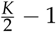 local minima, each inside an interval *I_i_*,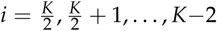. If *K* is odd, then the upper bound has 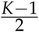 local minima, the first in interval 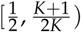, and each of the others in an interval *I_i_*, with 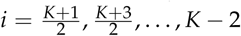. Note that because we restrict attention to *M* ∈ [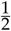,1), we do not count the point at *M* = 1 and *F* = 0 as a local minimum.

Theorem 2 from Appendix B describes the relative positions of the local minima within intervals *I_i_*, as a function of the number of subpopulations *K*. From Proposition 1 of Appendix B, for fixed *K*, the relative position of the local minimum within interval *I_i_* increases with *i*; as a result, the leftmost dips in the upper bound (those near *M* = 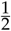) are less tilted toward the right endpoints of their associated intervals than are the subsequent dips (nearer *M* = 1). The unique local minimum in interval *I_i_* lies either exactly at 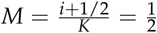 for the leftmost dip for odd *K* (Proposition 2)—or slightly to the right of the midpoint 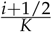 of interval *I_i_* in other intervals, but no farther from the center than 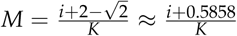 (Proposition 3).

The values of the upper bound on *F* at the local minima as a function of *i* are computed in Appendix B (eq. B.5) by substituting the positions *M* of the local minima into eq. 6. From Proposition 4 of Appendix B, for fixed *K*, the value of *F* at the local minimum in interval *I_i_* decreases as *i* increases. The maximal *F* among local minima increases as *K* increases (Proposition 5). The upper bound on *F* at the local minimum closest to *M* = 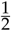 tends to 1 as K → ∞ (Proposition 5). The upper bound on *F* at the local minimum closest to M = 1, however, is always smaller than 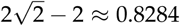 (Proposition 6).

In conclusion, although *F* is constrained below 1 for all values of *M* in the interior of intervals 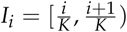, the constraint is reduced as *K* → ∞ and in the limit, it even completely disappears in the interval *I_i_*, closest to *M* = 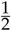. Nevertheless, there always exists a value of 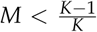 for which the upper bound on *F* is lower than 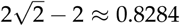.

### Mean range of possible F values

We now evaluate how strongly *M* constrains the range of *F* as a function of the number of subpopulations *K*. To do so, we compute the mean maximum *F* across all possible *M* values. This quantity, denoted *A*(*K*), corresponds to the area between the lower and upper bounds on *F* as a function of *M* divided by the length of the domain of possible *M* values, 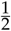. Small values of *A*(*K*) near 0 indicate a strong constraint, whereas large *A*(*K*) values near 1 indicate that for all *M*, *F* can range over most of the interval from 0 to 1. *A*(*K*) can also be interpreted as the mean maximum *F* attainable when *M* is uniformly distributed between 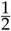 and 1.

Because the lower bound on *F* is 0 for all *M* between 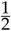 and 1, *A*(*K*) corresponds to the area under the upper bound on *F* divided by 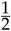, or twice the integral of eq. 6 between 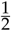 and 1:

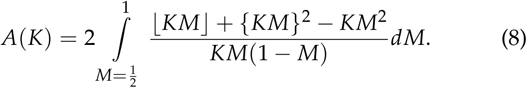

To compute *A*(*K*), we break the integral into a sum of integrals over intervals *I_i_*. If *K* is even, then we consider intervals 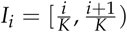, with 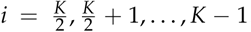. If *K* is odd, then we use the sum of integrals over intervals 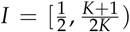 and 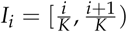, with 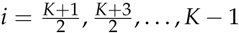.

By construction of *Q_i_*(*x*) (eq. 7), in each interval *I_i_*, the upper bound on *F* is equal to *Q_i_*(*x*), with *x* = {*KM*}. In the odd case, because 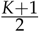 is an integer, on interval 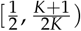, ⌊KM⌋ has a constant value 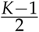 and 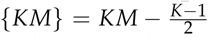, and the upper bound is equal to 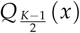. Making the substitution *x* = *KM* − *i*, we obtain *dx* = *K dM* and we can write eq. 8 in terms of *Q_i_*(*x*):

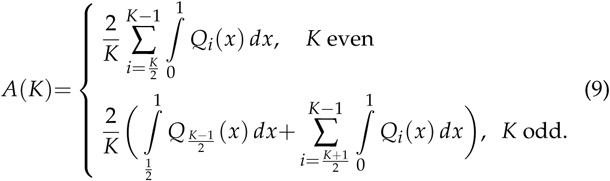

The integral is computed in Appendix C. We obtain, for both even and odd *K*:

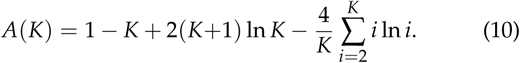

We also compute an asymptotic approximation *Ã*(*K*) of eq. 10 in Appendix C, producing the asymptotic relationship

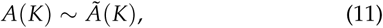

where

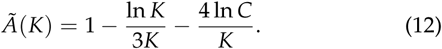

Here, *C* ≈ 1.2824 represents the Glaisher-Kinkelin constant.

For *K* = 2, *A*(*K*) = 2ln2 − 1 ≈ 0.3863, in accord with the *K* = 2 case of Jakobsson *et al.* (2013). Interestingly, the constraint on the mean range of *F* disappears as the number of subpopulations *K* → ∞. Indeed, from eqs. 11 and 12, we immediately see that lim_*K*→∞_ *A*(*K*) = 1 (Figure 2). As a mean of 1 indicates that *F* ranges from 0 to 1 for all values of *M* (except possibly on a set of measure 0), for large *K*, the range of *F* is approximately invariant with respect to the mean allele frequency *M*.

**Figure 2.**
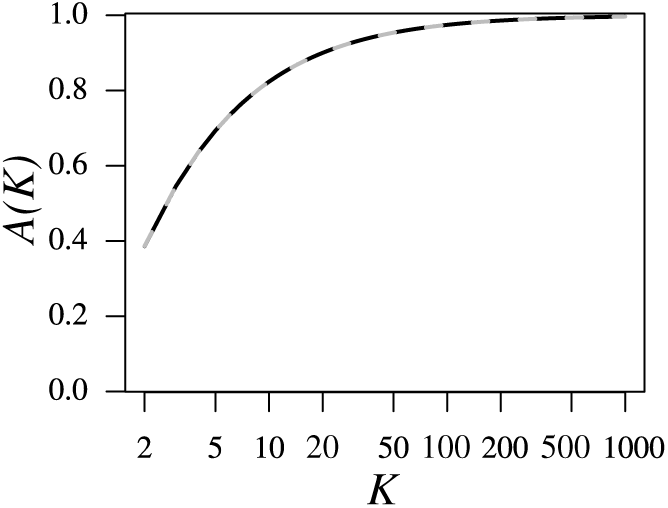
The mean *A*(*K*) of the upper bound on *F* over the interval *M* ∈ [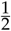,1), as a function of the number of subpopulations *K*. *A*(*K*) is computed from eq. 10 (black line). The approximation *Ã*(*K*) is computed from eq. 12 (gray dashed line). A numerical computation of the relative error of the approximation as a function of *K*, |*A*(*K*) − Ã (*K*)|/*A*(*K*), finds that the maximal error for 2 ≤ *K* ≤ 1000 is 0.00174, achieved when *K* = 2. The x-axis is plotted on a logarithmic scale.

The increase of *A*(*K*) with *K* is monotonic, as demonstrated in Theorem 3 from Appendix C. By numerically evaluating eq. 10, we find that although *A*(2) ≈ 0.3863, for *K* ≥ 7, *A*(*K*) exceeds 0.75, and for *K* ≥ 46, *A*(*K*) exceeds 0.95. Nevertheless, although the mean of the upper bound on *F* approaches 1, we have shown in Proposition 6 from Appendix B that for large *K*, values of *M* continue to exist at which the upper bound on *F* is constrained substantially below 1.

## Evolutionary processes and the joint distribution of *M* and *F* for a biallelic marker and *K* subpopulations

To illustrate the mathematical properties of *F* in the context of evolutionary models, we simulated the joint distribution of *F* and *M* under simple biological scenarios, and compared this distribution to the mathematical bounds on *F*. This analysis considers allele frequency distributions generated by evolutionary models, rather than treating *M* as uniformly distributed in [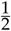,1).

### Simulations

We simulated independent single-nucleotide polymorphisms under the coalescent, using the software MS (Hudson 2002). We considered a population of total size *KN* diploid individuals subdivided into *K* subpopulations of equal size *N*. At each generation, a proportion *m* of the individuals in a subpopulation originated from another subpopulation, with the subpopulations of origin determined by the migration model. Thus, the scaled migration rate is 4*Nm*, and it corresponds to twice the number of individuals in a subpopulation that originate elsewhere.

We considered three migration models (Figure 3), the finite island model (Maruyama 1970; Wakeley 1998) and the finite rectangular and linear stepping-stone models (Maruyama 1970; Wilkinson-Herbots 1998). In the island model, migrants have the same probability 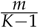 to come from any other subpopulation. In the rectangular stepping-stone model, subpopulations are arranged on a rectangular bounded habitat. Each subpopulation receives migrants from each adjacent subpopulation with the same probability. Subpopulations not on the habitat boundaries receive migrants at the same rate 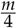 from each of four adjacent subpopulations; subpopulations on habitat edges receive migrants from each of three adjacent subpopulations at rate 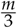; subpopulations at vertices receive migrants from each of two adjacent subpopulations at rate 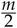. In the linear stepping-stone model, subpopulations are arranged along a linear bounded habitat. Each subpopulation receives migrants from each adjacent subpopulation at the same rate. We consider reflecting boundaries, so that interior subpopulations receive migrants at rate 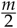 from each of two adjacent subpopulations, whereas subpopulations at habitat boundaries receive migrants from a single adjacent subpopulation at rate *m*.

**Figure 3.**
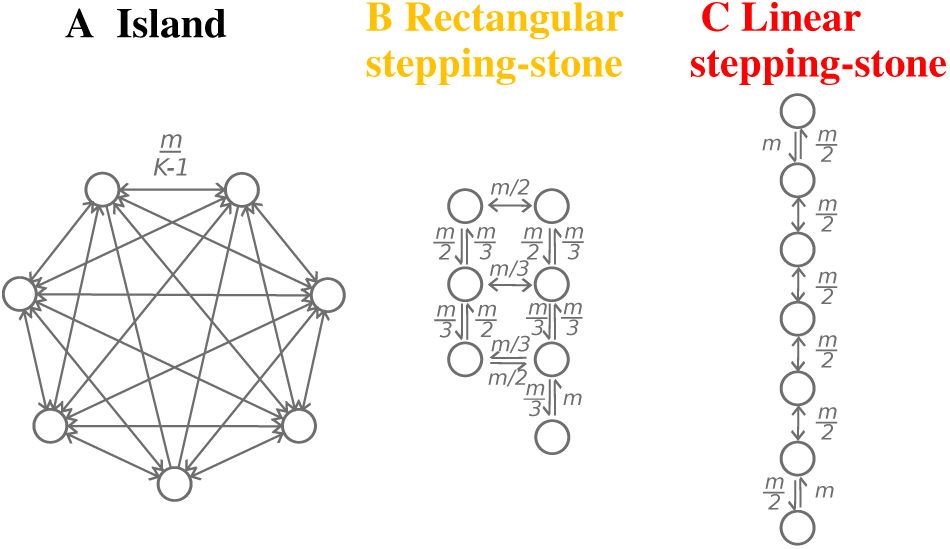
Three migration models. (A) Island model. (B) Rectangular stepping-stone model. (C) Linear stepping-stone model. Quantities on the arrows represent the backward migration rates between pairs of subpopulations.

We examined three values of *K*: 2, 7, and 40. For *K* = 2, all three migration models are equivalent. Under the rectangular stepping-stone model, for *K* = 7, we considered a habitat of 4 × 2 subpopulations with one subpopulation missing at the edge (Figure 3B); for *K* = 40, we considered an 8 × 5 habitat. We used three values of 4*Nm*: 0.1, 1, and 10. Note that under the coalescent model in MS, time is scaled in units of 4*N* generations, so there is no need to specify the subpopulation sizes *N*. To obtain independent SNPs, we used the MS command -s to fix the number of segregating sites to 1. For each parameter pair (*K*, 4*Nm*), we performed 100,000 replicate simulations, sampling 100 sequences per subpopulation in each replicate. *F* values were computed from the parametric allele frequencies.

Fixing the number of segregating sites to 1 and accepting all coalescent genealogies entails an implicit assumption that all genealogies have equal potential to produce exactly one segregating site. We therefore also considered a different approach to generating SNPs; instead of fixing the number of SNPs to 1, we assumed an infinitely-many-sites model with a specified scaled mutation rate *θ*, discarding simulations leading to more than 1 segregating site. We chose *θ* so that the expected number of segregating sites in a constant-sized population—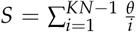— was 1. This approach produces similar results to the fixed-*S* simulation (Figure S1). MS commands appear in File S1.

### Weak migration in the island model

Under the island model with weak migration (4*Nm* = 0.1), the joint distribution of *M* and *F* is highest near the upper bound on *F* in terms of *M*, for all *K* (Figure 4A-C). For *K* = 2, most SNPs have *M* near 0.5, representing fixation of the major allele in one subpopulation and absence in the other, and *F* near 1 (orange and red areas in Figure 4A). The mean *F* in sliding windows for *M* closely follows the upper bound on *F* in terms of *M*. For *K* = 7, most SNPs have *M* near 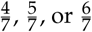, representing fixation of the major allele in 4, 5, or 6 subpopulations and absence in the other subpopulations, and *F* ≈ 1 (Figure 4B). The mean *F* closely follows the upper bound on *F*. For *K* = 40, most SNPs either have *M* near 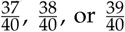, and *F* ≈ 1, or 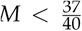 and *F* ≈ 0.92 (Figure 4C). The mean *F* follows the upper bound on *F* for 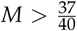. For 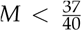, it lies below the upper bound and does not possess the peaks characteristic of the upper bound.

**Figure 4.**
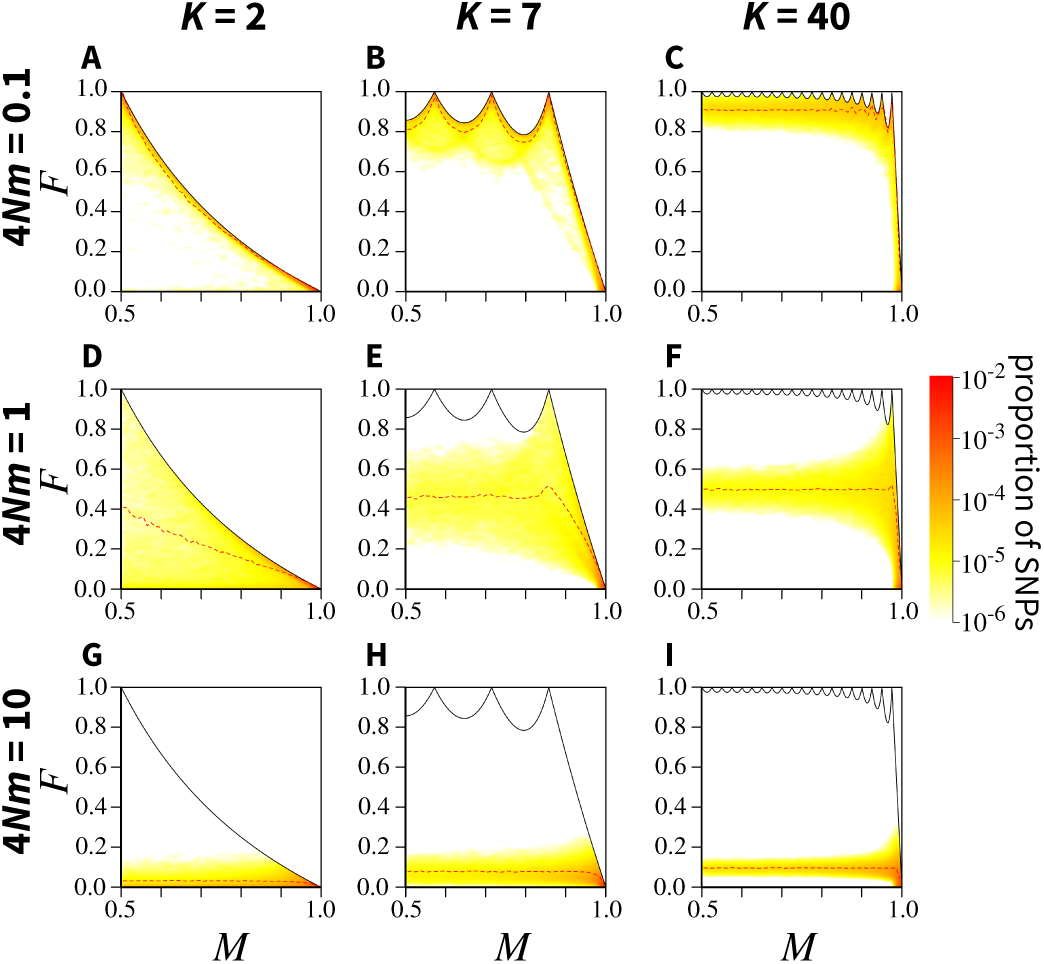
Joint density of the frequency *M* of the most frequent allele and *F* in the island model, for different numbers of sub-populations *K* and scaled migration rates 4*Nm* (where *N* is the subpopulation size and m the migration rate). (A) *K* = 2, 4*Nm* = 0.1. (B) *K* = 7, 4*Nm* = 0.1. (C) *K* = 40, 4*Nm* = 0.1. (D) *K* = 2, 4*Nm* = 1. (E) *K* = 7, 4*Nm* = 1. (F) *K* = 40, 4*Nm* = 1. (G) *K* = 2, 4*Nm* = 10. (H) *K* = 7, 4*Nm* = 10. (I) *K* = 40, 4*Nm* = 10. The black solid line represents the upper bound on *F* in terms of *M* (eq. 6); the red dashed line represents the mean *F* in sliding windows of *M* of size 0.02 (plotted from 0.51 to 0.99). Colors represent the density of SNPs, estimated using a Gaussian kernel density estimate with a bandwidth of 0.007, with density set to 0 outside of the bounds. SNPs are simulated using coalescent software MS, assuming an island model of migration and conditioning on 1 segregating site. See Figure S1 for an alternative algorithm to simulating SNPs. Each panel considers 100,000 replicate simulations, with 100 lineages sampled per subpopulation.

Coalescent theory provides a framework to understand these observations. Wakeley (1999) showed that in the limit in which the migration rate is much lower than the coalescence rate (i.e., 4*Nm* ≪ 1), coalescence follows two phases. In the scattering phase, lineages coalesce in each subpopulation, leading to a state with a single lineage per subpopulation. In the collecting phase, lineages from different subpopulations coalesce. As a result, considering *K* subpopulations with equal sample size *n*, when 4*Nm* ≪ 1, genealogies tend to have *K* long branches close to the root, each corresponding to a subpopulation and each leading to n shorter terminal branches. These *K* long branches coalesce as pairs of them accumulate by migration in shared ancestral subpopulations. A random mutation on such a genealogy is likely to occur in one of two places. It can occur on a long branch during the collecting phase, in which case the derived allele will have a frequency of 1 in all subpopulations whose lineages descend from the branch, and 0 in the other subpopulations. Alternatively, it can occur toward the terminal branches in the scattering phase, in which case the mutation will be at frequency *p_k_* > 0 in a single subpopulation, and at frequency 0 in all others. These scenarios that are likely under weak migration—in which one of the alleles is fixed in some subpopulations or present only in a single subpopulation—correspond closely to conditions under which the upper bound on *F* is reached at fixed *M*. Thus, the properties of likely genealogies under weak migration explain the proximity of *F* to its upper bound.

### Intermediate migration in the island model

Under the island model with intermediate migration (4*Nm* = 1), for all *K*, the joint density of *M* and *F* is highest at lower values of *F* than in the case of weak migration (Figure 4D-F). For *K* = 2, most SNPs have *M* > 0.8, and the mean *F* in sliding windows for *M* is almost equidistant from the upper and lower bounds on *F* in terms of *M*, nearing the upper bound as *M* increases (Figure 4D). For *K* = 7, most SNPs have *M* > 0.9; as was seen for *K* = 2, the mean *F* in sliding windows for *M* is almost equidistant from the upper and lower bounds on *F*, moving toward the upper bound as *M* increases (Figure 4E). For *K* = 40, the pattern is comparable, most SNPs having *M* > 0.95 (Figure 4F).

With intermediate migration, migration is sufficient that more mutations than in the weak-migration case generate polymorphism in multiple subpopulations. A random mutation is likely to occur on a branch that leads to many terminal branches from the same subpopulation, but also to branches from other subpopulations. Thus, the allele is likely to have intermediate frequency in multiple subpopulations. This configuration does not generate the conditions under which the upper bound on *F* is reached, so that except at the largest *M*, intermediate migration leads to values that are not as close to the upper bound as in the weak-migration case. For large *M*, the rarer allele is likely to be only in one subpopulation, so that *F* is nearer to the upper bound.

### Strong migration in the island model

With strong migration (4*Nm* = 10), the joint density of *M* and *F* nears the lower bound on *F* in terms of *M* (Figure 4G-I). For each choice of *K*, most SNPs have *M* > 0.9 and *F* ≈ 0, with the mean *F* increasing somewhat as *K* increases from 2 to 7 and 40.

In the limit in which migration is strong, because lineages can migrate between subpopulations quickly, they can also coalesce quickly, irrespective of their subpopulations of origin. As a result, a random mutation is likely to occur on a branch that leads to terminal branches from many subpopulations. The allele is expected to be at comparable frequency in all subpopulations, so that *F* is likely to be small. This scenario corresponds to the conditions under which the lower bound on *F* is approached.

### Rectangular and linear stepping-stone models

Under the rectangular stepping-stone model, properties of *F* in relation to *M* are qualitatively similar to those under the island model, but with higher *F* (Figures 5B,E,H and 6B,E,H). For a fixed number of subpopulations *K*, the geometry in the rectangular stepping-stone model, with 2 to 4 connections per subpopulation, generates less migration among the subpopulations, so that the genetic difference among subpopulations is higher than in the fully connected graph of the island model. Thus, with *M*, *K*, and 4*Nm* held constant, *F* is generally higher in the rectangular stepping-stone model.

**Figure 5.**
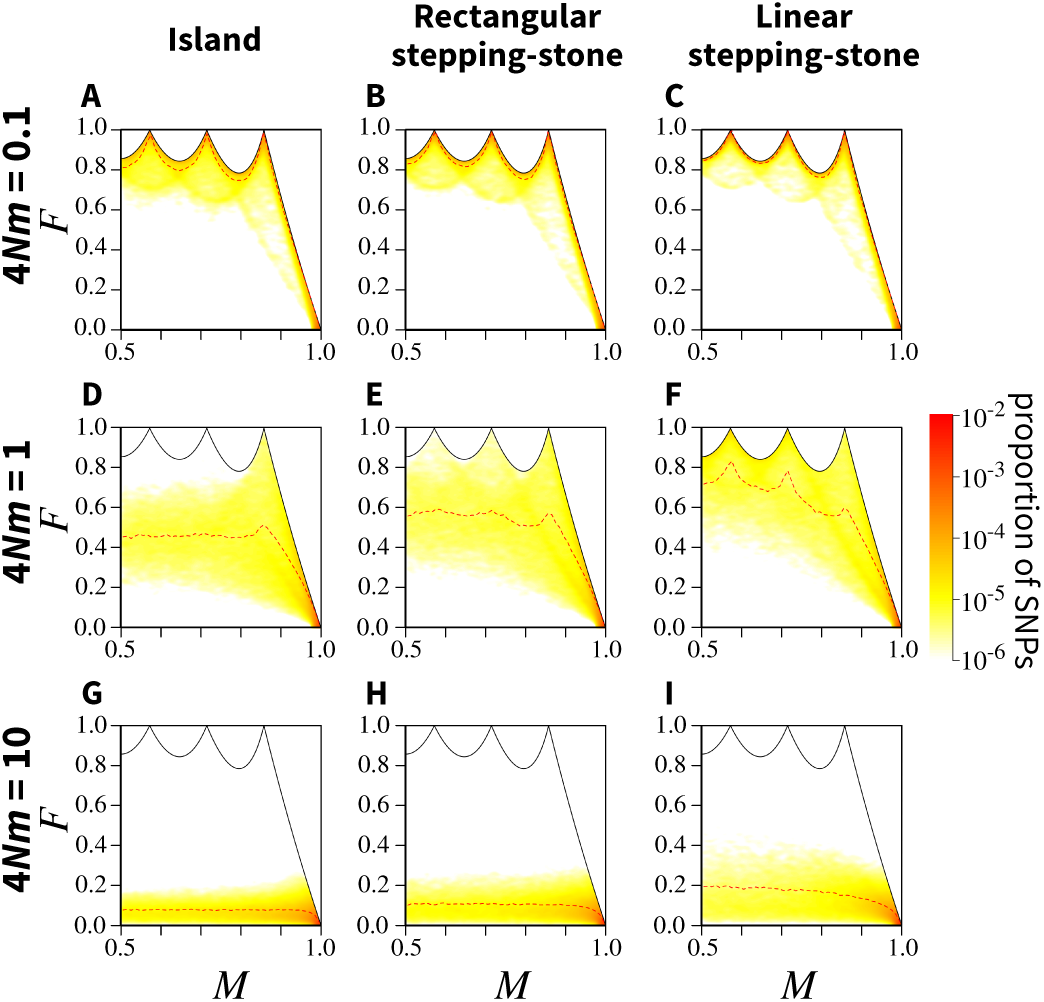
Joint density of the frequency *M* of the most frequent allele and *F*, for different migration models and scaled migration rates 4*Nm*, considering *K* = 7 subpopulations. Panels A,D,G for the island model are copied from Figure 4B,E,H for ease of comparison. The simulation procedure and figure design follow Figure 4.

**Figure 6.**
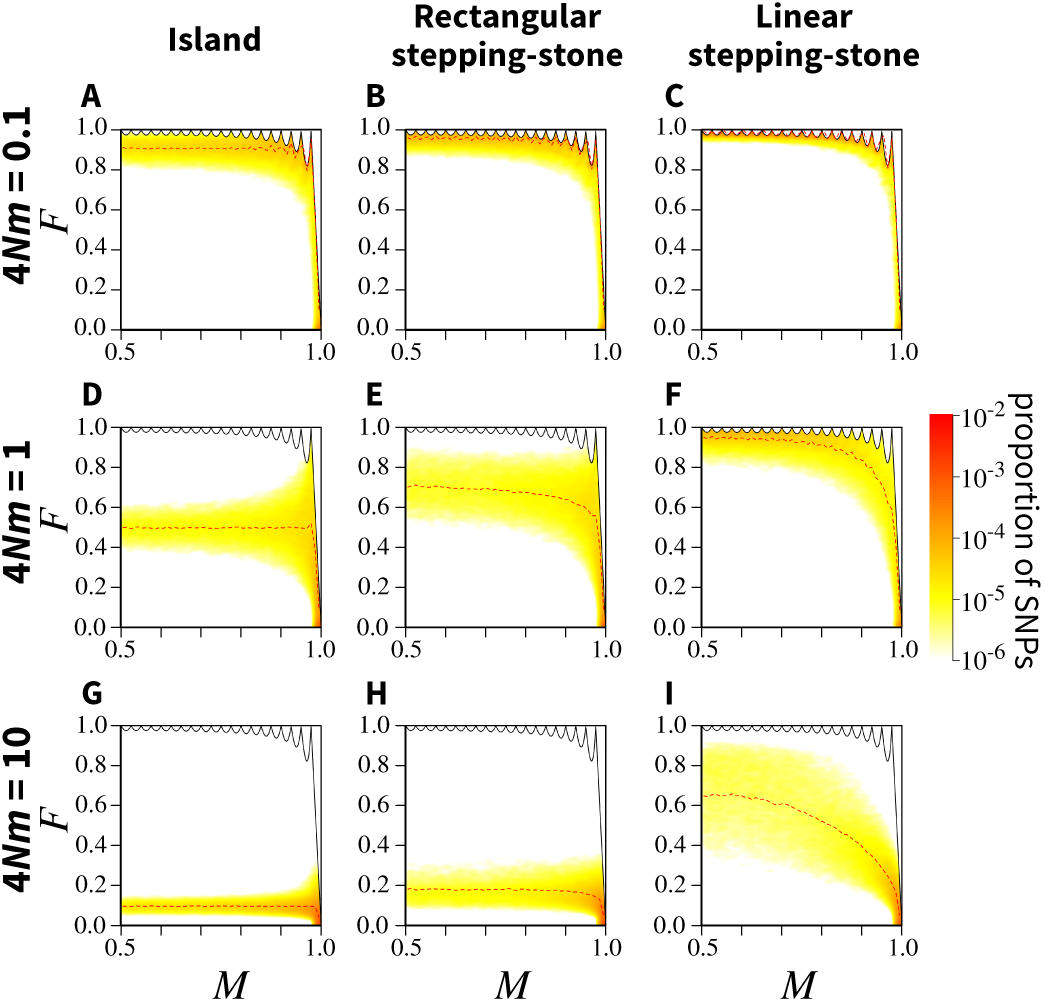
Joint density of the frequency *M* of the most frequent allele and *F*, for different migration models and scaled migration rates 4*Nm*, considering *K* = 40 subpopulations. Panels A,D,G for the island model are copied from Figure 4C,F,I for ease of comparison. The simulation procedure and figure design follow Figure 4.

In the linear stepping-stone model, *F* is higher still than in the rectangular model (Figures 5C,F,I and 6C,F,I). Connectivity among subpopulations is reduced, with each subpopulation having only 1 or 2 neighbors. The probability that a mutation remains localized and fixed in some subpopulations while being absent in others is greater than in the other geometries, so that *F* exceeds that observed in the other models.

### Proximity of the joint density of M and F to the upper bound

To illustrate the influence of evolutionary processes on the relationship of *F* to the upper bound and to summarize the features of Figures 4, 5, and 6, we can quantify the proximity of the joint density of *M* and *F* to the bounds on *F* in terms of *M*, as a function of *K*, 4*Nm* and the migration models.

For a set of *Z* loci, denote by *F_z_* and *M_z_* the values of *F* and *M* at locus *z*, respectively. The mean *F* for the set, denoted *F̅*, is

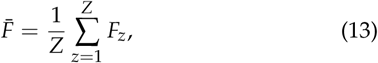

A corresponding mean maximum *F* given the observed *M_z_*, *z* = 1,2,…, *Z*, denoted *F*_max_, is the sum of the maximal values of *F* across the *Z* loci (eq. 6):

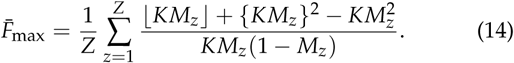

The ratio *F̅* / *F̅*_max_ gives a sense of the proximity of the *F* values to their upper bounds: it ranges from 0, when *F* values at all SNPs equal their lower bounds, to 1, when *F* values at all SNPs equal their upper bounds. Figure 7 shows the ratio *F̅* /*F̅*_max_ under the three migration models, for different values of *K* and 4*Nm*.

**Figure 7.**
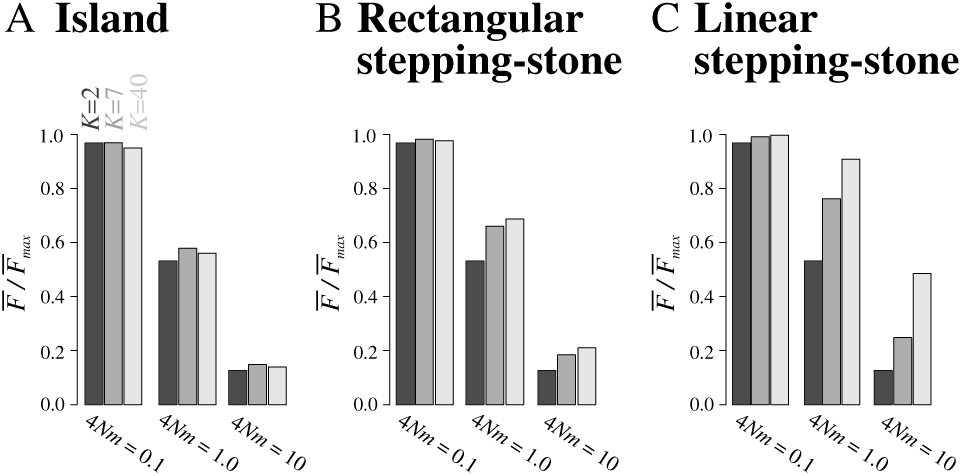
*F̅*/F̅_max_, the ratio of the the mean *F* to the mean maximal *F* given the observed frequency of the most frequent allele *M*, as a function of the number of subpopulations *K* and the scaled migration rate 4*Nm*, for three migration models. (a) Island model. (b) Rectangular stepping-stone model. (c) Linear stepping-stone model. Colors represent values of *K*: 2, 7, and 40. *F* values are computed from coalescent simulations using MS, for 10,000 independent SNPs and 100 lineages sampled per subpopulation. *F̅*_max_ is computed from eq. 14.

For each model and each value of the number of subpopulations, *F̅*/*F̅*_max_ decreases with 4*Nm*. This result summarizes the influence of the migration rate observed in Figures 4, 5, and 6: *F* values tend to be close to the upper bound under weak migration, and near the lower bound under strong migration.

For a fixed number of subpopulations and a fixed scaled migration rate, *F̅*/*F̅_max_* is smaller under the island model than under the rectangular stepping-stone model, and smaller under the rectangular model than under the linear model. This observation can be explained by the stronger constraints on migration in the linear case, in which immigrants come from at most 2 other subpopulations, than in the rectangular case, with up to 4 neighbors, and the island model, with *K* − 1. The smaller number of neighbors prevents genetic homogenization between subpopulations and thus leads to larger *F* values.

Under the island model, *F̅*/*F̅*_max_ is only minimally influenced by the number of subpopulations *K* (Figure 7A). Even though the upper bound on *F* in terms of *M* is strongly affected by *K*, the proximity of *F* to the upper bound is similar across *K* values. Under the rectangular and linear stepping-stone models, however, *K* has a stronger influence on *F̅*/ *F̅_max_*, which increases with *K* (Figure 7B,C). This result can be explained by noting that unlike in the island model, which is fully connected irrespective of the number of subpopulations, at a fixed migration rate, the increasing number of subpopulations produces greater isolation of distant subpopulations in the stepping-stone models, generating greater genetic differentiation and thus leading to *F* values closer to their upper bounds.

## Application to human genomic data

We now use our theoretical results to explain observed patterns of human genetic differentiation, and in particular, to explain the impact of the number of subpopulations. We employ data from Li *et al.* (2008) on 577,489 SNPs from 938 individuals of the Human Genome Diversity Panel (HGDP; Cann *et al.* 2002), as compiled by Pemberton *et al.* (2012). We use the same division of the individuals into seven geographic regions that was examined by Li *et al.* (2008) (Africa, Middle East, Europe, Central and South Asia, East Asia, Oceania, America). We computed the parametric allele frequencies for each region, averaging across regions to obtain the frequency *M* of the most frequent allele. We then computed *F* for each SNP, averaging F values across SNPs to obtain the overall *F* for the full SNP set.

To assess the impact of the number of subpopulations *K* on the relationship between *M* and *F*, we computed *F* for all 120 sets of two or more geographic regions (Figure 8). The 21 pairwise *F* values range from 0.007 (between Middle East and Europe) to 0.101 (Africa and America), with a mean of 0.057, standard deviation of 0.027, and median of 0.061. *F* is substantially larger for sets of three geographic regions. The smallest value is larger, 0.012 (Middle East, Europe, Central/South Asia), as is the largest value, 0.133 (Africa, Oceania, America), the mean of 0.076, and the median of 0.089. Among the 21 × 5 = 105 ways of adding a third region to a pair of regions, 83 produce an increase in *F*. For 17 sets of three regions, the value of *F* exceeds that for each of its three component pairs.

**Figure 8.**
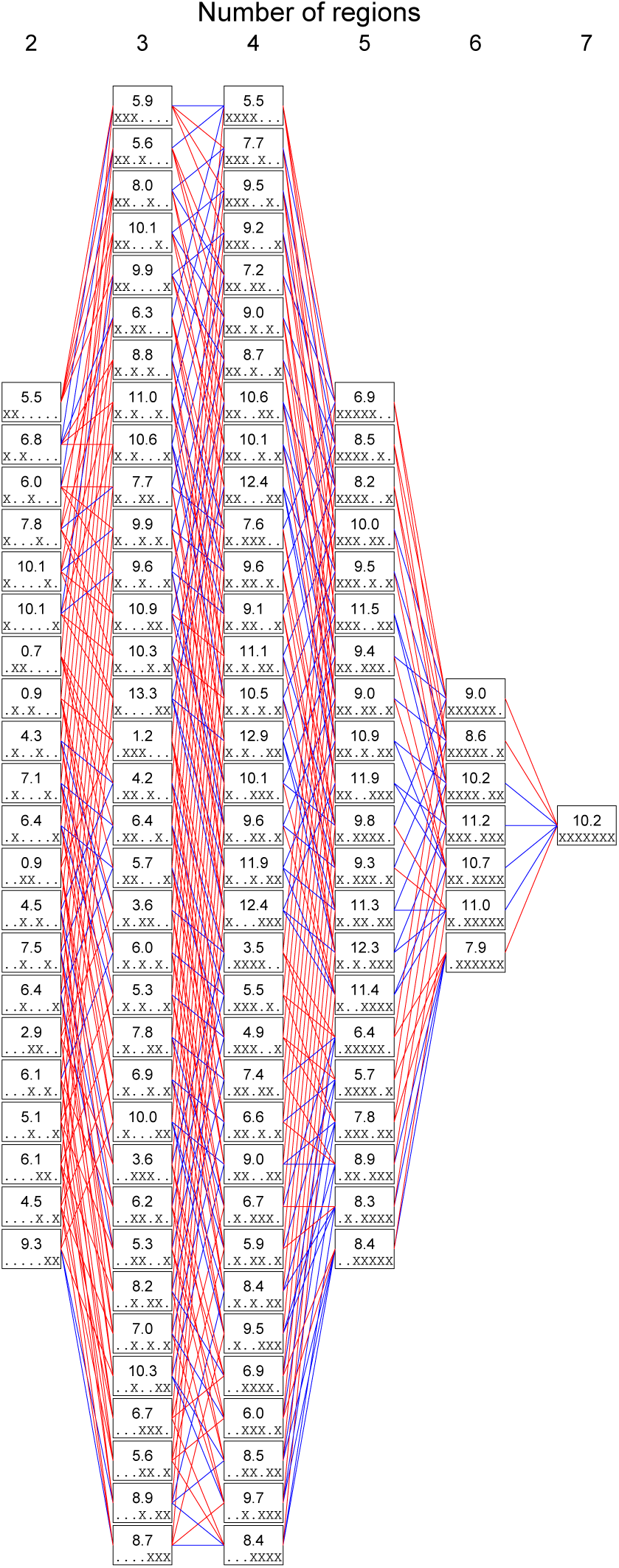
Mean *F* values across loci for sets of geographic regions. Each box represents a particular combination of 2, 3, 4, 5, 6, or all 7 geographic regions. Within a box, the numerical value shown is 100 × *F* among the regions. The regions considered are indicated by the pattern of “.” and “X” symbols within the box, with “X” indicating inclusion and “.” indicating exclusion. From left to right, the regions are Africa, Middle East, Europe, Central/South Asia, East Asia, Oceania, America. Thus, for example, X…X. indicates the subset {Africa, East Asia}. Lines are drawn between boxes that represent nested subsets. A line is colored red if the larger subset has a higher *F* value, and blue if it has a lower *F*. Computations rely on 577,489 SNPs from the Human Genome Diversity Panel.

The pattern of increase of *F* with the inclusion of additional subpopulations can be seen in Figure 9A, which plots the *F* values from Figure 8 as a function of *K*. The magnitude of the increase is greatest from *K* = 2 to *K* = 3, decreasing with increasing *K*. From *K* = 3 to 4, 82 of 140 additions of a region increase *F*; 54 of 105 produce an increase from *K* = 4 to 5, 21 of 42 from *K* = 5 to 6, and 3 of 7 from *K* = 6 to 7. The seven-region *F* of 0.102 exceeds all the pairwise *F* values.

**Figure 9.**
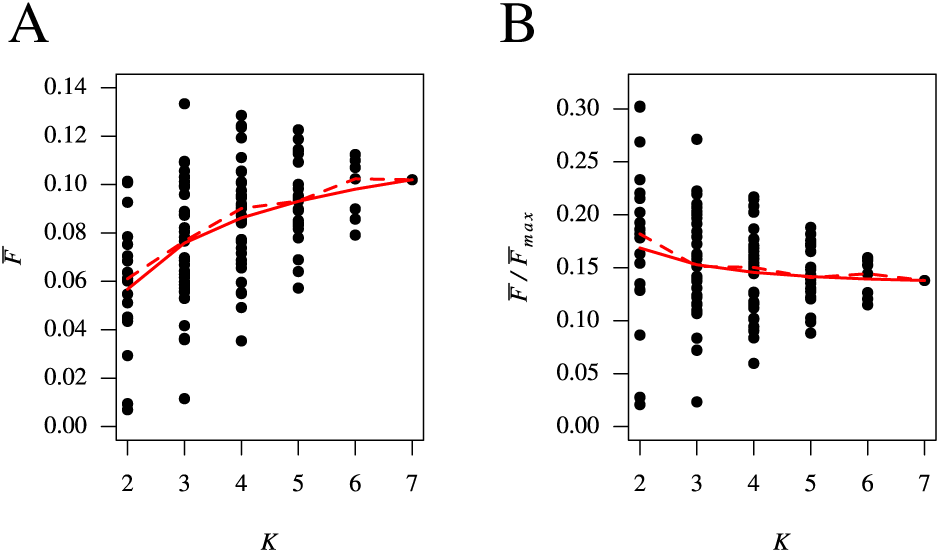
*F* values for sets of geographic regions as a function of *K*, the number of regions considered. (A) *F̅*, computed using eq. 13. (B) *F̅*/*F̅*_max_, computed using eq. 14. For each subset of populations, the value of *F* is taken from Figure 8. The mean across subsets for a fixed *K* appears as a solid red line, and the median as a dashed red line.

The larger *F* values with increasing *K* can be explained by the difference in constraints on *F* in terms of *M* (Figure 10). For fixed *M*, as we saw in the increase of *A* (*K*) with *K* (Figure 2), the permissible range of *F* values is smaller on average for *F* values computed among smaller sets of populations than among larger sets. For example, the maximal *F* value at the mean *M* of 0.76 observed in pairwise comparisons is 0.33 for *K* = 2 (black line in Figure 10A), while the maximal *F* value at the mean *M* of 0.77 observed for the global comparison of seven regions is 0.86 for *K* = 7 (Figure 10B). Given the stronger constraint in pairwise calculations, it is not unexpected that pairwise *F* values would be smaller than the values computed with more regions, such as in the full 7-region computation. Interestingly, the effect of *K* on *F* is largely eliminated when *F* values are normalized by their maxima (Figure 9B). The normalization, which takes both *K* and *M* into account, generates a nearly constant trend in the mean and median of *F* as a function of *K*, with higher values for *K* = 2.

**Figure 10.**
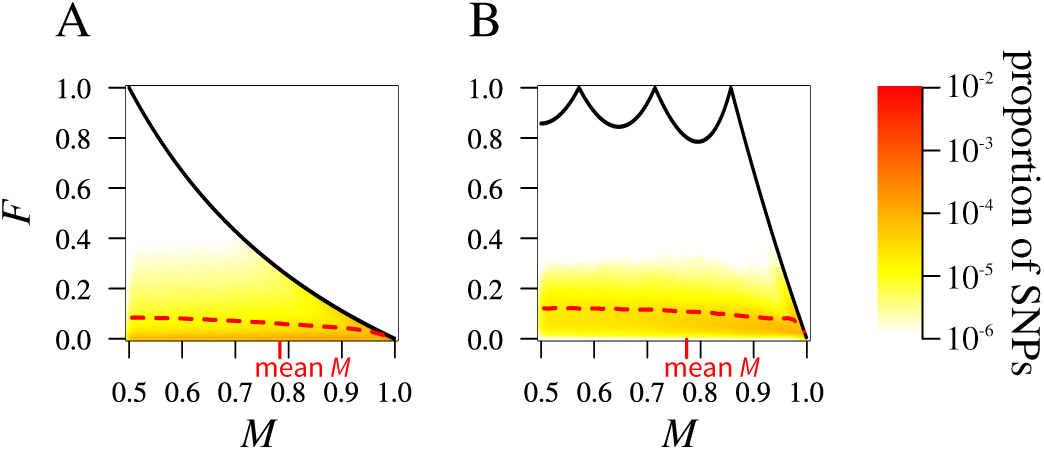
Joint density of the frequency *M* of the most frequent allele and *F* in human population-genetic data, considering 577,489 SNPs. (A) *F* computed for pairs of geographic regions. The density is computed from the set of *F* values for all 21 pairs of regions. (B) *F* computed among *K* = 7 geographic regions. The figure design follows Figure 4.

## Discussion

We have evaluated the constraint imposed by the frequency *M* of the most frequent allele on the range of *F_ST_*, for arbitrary many subpopulations. Although the range of *F_ST_* is unconstrained within the unit interval for a finite set of values of 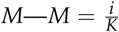, where *i* is an integer greater than or equal to 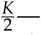it is constrained below 1 for all other values of *M*. We have found that the number of subpopulations *K* has a considerable impact on the range of *F_ST_*, with a weaker constraint on *F_ST_* as *K* increases. As was shown by Jakobsson *et al.* (2013), for *K* = 2, considering all possible values of *M*, *F_ST_* values are restricted to 38.63% of the possible space. Considering *K* = 100, however, *F_ST_* values can occupy 97.47% of the space. Although the mean over *M* values of the permissible interval for *F_ST_* approaches the full unit interval as *K* → ∞, we find that for any *K*, there exists an allele frequency *M* < (*K* − 1) /*K* for which the maximal *F* is lower than 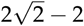.

Multiple studies have highlighted the relationship between *F_ST_* and *M* in two subpopulations, for biallelic markers (Rosenberg *et al.* 2003; Maruki *et al.* 2012), and more generally, for an unspecified (Jakobsson *et al.* 2013) or specified number of alleles (Edge and Rosenberg 2014). We have extended these results to the case of biallelic markers in a specified but arbitrary number of subpopulations, comprehensively describing the relationship between *F_ST_* and *M* for the biallelic case. The study is part of an increasing body of work characterizing the mathematical relationship of population-genetic statistics with quantities that constrain them (Hedrick 1999, 2005; Rosenberg and Jakobsson 2008; Reddy and Rosenberg 2012). As we have seen, such relationships contribute to understanding the behavior of the statistics in evolutionary models and to interpreting counterintuitive results in human population genetics.

### Properties of *F_ST_* in evolutionary models

Our work extends classical results about the impact of evolutionary processes on *F_ST_* values. Wright (1951) showed that in an equilibrium population, *F_ST_* is expected to be near 1 if migration is weak, and near 0 if migration is strong. On the basis of our simulations, we can more precisely formulate this proposition: considering a SNP *at frequency M* in an equilibrium population, *F_ST_* is expected to be *near its upper bound in terms of *M* if migration is weak* and near 0 if migration is strong. This formulation of Wright’s proposition makes it possible to explain why SNPs subject to the same migration process can display a variety of *F_ST_* patterns; indeed, under weak migration, we expect *F_ST_* values to mirror the considerable variation in the upper bound on *F_ST_* in terms of *M*.

We also provide a framework for interpreting properties of *F_ST_* across multiple migration models. Maruyama (1970) showed that under the linear stepping-stone model, *F_ST_* values tend to be closer to 1 than under an island model. Our results provide a more precise formulation of this classical pattern: under stepping-stone models, *F_ST_* values tend to be closer to their upper bound in terms of *M* than under an island model.

### Lower *F_ST_* values in pairwise comparisons than in comparisons of more subpopulations

*F_ST_* values have often been compared across computations with different numbers of subpopulations. Such comparisons appear frequently, for example, in studies of domesticated animals such as horses, pigs, and sheep (Cañon *et al.* 2000; Kim *et al.* 2005; Lawson Handley *et al.* 2007). In human populations, Table 1 of the microsatellite study of Rosenberg *et al.* (2002) presents comparisons of *F_ST_* values for scenarios with *K* ranging from 2 to 52. Table 3 of Rosenberg *et al.* (2006) compares *F_ST_* values for microsatellites and biallelic indels in population sets with *K* ranging from 2 to 18. Major SNP studies have also compared *F_ST_* values for scenarios with *K* = 2 and *K* = 3 groups (Hinds *et al.* 2005; International HapMap Consortium 2005).

Our results suggest that such comparisons between *F_ST_* values with different *K* can hide an effect of the number of subpopulations, especially when some of the comparisons involve the most strongly constrained case of *K* = 2. For human genetic differentiation, we found that owing to a difference in the *F_ST_* constraint for different *K* values, pairwise *F_ST_* values between continental regions were consistently lower than *F_ST_* computed using three or more regions, and sets of three regions were identified for which the *F_ST_* value exceeded the values for all three pairs of regions in the set. The effect of *K* might help illuminate why SNP-based pairwise *F_ST_* values between human populations (Table S11 of 1000 Genomes Project Consortium *et al.* 2012) are generally smaller than estimates that use all populations together (11.1% of genetic variance due to between-region or between-population differences; Li *et al.* 2008). Our results suggest that comparing *F_ST_* values with different choices of *K* can generate as much difference—twofold—as comparing *F_ST_* with different marker types (Holsinger and Weir 2009). This substantial impact of *K* on *F_ST_* merits further attention.

### Consequences for the use of *F_ST_* as a test statistic

The effects of constraints on *F_ST_* extend beyond the use of *F_ST_* as a statistic for genetic differentiation. In *F_ST_*-based genome scans for local adaptation, tracing to the work of Lewontin and Krakauer (1973), a hypothesis of spatially divergent selection at a candidate locus is evaluated by comparing *F_ST_* at the locus with the *F_ST_* distribution estimated from a set of putatively neutral loci. Under this test, *F_ST_* values smaller or larger than expected by chance are interpreted as being under stabilizing or divergent selection, respectively. Modern versions of this approach compare *F_ST_* values at single loci with the distribution across the genome (Beaumont and Nichols 1996; Akey *et al.* 2002; Foll and Gaggiotti 2008; Bonhomme *et al.* 2010; Günther and Coop 2013).

The constraints on *F_ST_* in our work and the work of Jakobsson *et al.* (2013) and Edge and Rosenberg (2014) suggest that *F_ST_* values strongly depend on the frequency of the most frequent allele. Consequently, we expect that *F_ST_* outlier tests that do not explicitly take into account this constraint will result in a deficit of power at loci with high- and low-frequency alleles. Because pairwise *F_ST_* and *F_ST_* values in many populations have different constraints, we predict that the effect of the constraint on outlier tests relying on a single global *F_ST_* (e.g., Beaumont and Nichols 1996; Foll and Gaggiotti 2008) will be smaller than in tests relying on pairwise *F_ST_* (e.g., Günther and Coop 2013).

### Conclusion

Many recent articles have noted that *F_ST_* often behaves counterintuitively (Whitlock 2011; Alcala *et al.* 2014; Wang 2015)—for example, indicating low differentiation in cases in which populations do not share any alleles (Balloux *et al.* 2000; Jost 2008) or suggesting less divergence among populations than is visible in clustering analyses (Tishkoff *et al.* 2009; Algee-Hewitt *et al.* 2016). It has thus become clear that observed *F_ST_* patterns often trace to peculiar mathematical properties of *F_ST_*—in particular its relationship to other statistics such as homozygosity or allele frequency—instead of to biological phenomena of interest. Our work here, extending approaches of Jakobsson *et al.* (2013) and Edge and Rosenberg (2014), seeks to characterize those properties, so that the influence of mathematical constraints on *F_ST_* can be disentangled from biological phenomena.

In response to a mathematical dependence of *F_ST_* on the within-subpopulation mean heterozygosity *H_S_*, Wang (2015) has proposed plotting the joint distribution of *H_S_* and *F_ST_*, in order to assess the correlation between the two statistics. Using the island model, Wang (2015) argued that when *H_S_* and *F_ST_* are uncorrelated, *F_ST_* is expected to be more informative about the demographic history of a species than when they are strongly correlated and *F_ST_* is merely a reflection of the within-subpopulation diversity. Our results suggest a related framework: studies can compare plots of the joint distribution of *M* and *F_ST_* with the bounds on *F_ST_* in terms of *M*. This framework, which examines constraints on *F_ST_* in terms of allele frequencies in the total population, complements that of Wang (2015), which considers constraints in terms of subpopulation allele frequencies. Such analyses, considering *F_ST_* together with additional measures of allele frequencies, are desirable in diverse scenarios—for explaining counterintuitive *F_ST_* phenomena, for avoiding overinterpretation of *F_ST_* values, and for making sense of *F_ST_* comparisons across settings that have a substantial difference in the nature of one or more underlying parameters.

## Acknowledgements.

We acknowledge support from NSF grants BCS-1515127 and DBI-1458059, from NIJ grant 2014-DN-BX-K015, a postdoctoral fellowship from the Stanford Center for Computational, Evolutionary, and Human Genomics, and a Swiss National Science Foundation Early Postdoc.Mobility fellowship.

## Appendix A. Demonstration of eq. 5

This appendix provides the derivation of the upper bound on 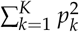 as a function of *K* and *M*.

### Theorem 1.

*Suppose σ* > 0 *and K* ≥ ⌊*σ*⌋ + 1 *are specified, where K is an integer. Considering all sequences* 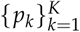 *with p_k_* ∈ [0,1], 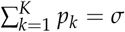, *and k* < ℓ *implies p_k_* < *p*_ℓ_, 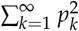 *is maximal if and only if p_k_* = 1 *for* 1 ≤ *k* ≤ ⌊*σ*⌋, *p*_⌊*σ*⌋+1_ = *σ* − ⌊*σ*⌋, *and p_k_* = 0 *for k* > ⌊*σ*⌋ + 1, *and its maximum is* (*σ* − ⌊*σ*⌋)^2^ + ⌊*σ*⌋.

*Proof.* This theorem is a special case of Lemma 3 from Rosenberg and Jakobsson (2008), which states (changing notation for some of the variables to avoid confusion): “Suppose *A* > 0 and *C* > 0 and that ⌈*C*/*A*⌉ is denoted *L*. Considering all sequences 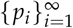 with 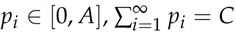, and *i* < *j* implies *p_i_* ≥ *p_j_*, 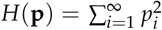 is maximal if and only if *p_i_*, = *A* for 1 ≤ *i* ≤ *L* − 1, *p_L_* = *C* − ( *L* − 1) *A*, and *p_i_* = 0 for *i* > *L*, and its maximum is *L*(*L* − 1) *A*^2^ − 2*C*(*L* − 1) *A* + *C*^2^.”

In our special case, we apply the lemma with *A* = 1 and *C* = *σ*. We also restrict consideration to sequences of finite rather than infinite length; however, our condition *K* ≥ ⌊*σ*⌋ + 1 for the number of terms in the sequence guarantees that the maximum in the case of infinite sequences—which requires ⌈*σ*⌉ ≤ ⌊*σ*⌋ + 1 nonzero terms—is attainable with sequences of the finite length we consider. For convenience in numerical computations, we state our result using the floor function rather than the ceiling function, requiring some bookkeeping to obtain our corollary.

If *σ* is not an integer, then in Lemma 3 of Rosenberg and Jakobsson (2008), *L* = ⌊*σ*⌋ + 1, and the maximum occurs with *p*_1_ = *p*_2_ = … = *p*_⌊*σ*⌋_ = 1, *p*_⌊*σ*⌋+1_ = *σ* − ⌊*σ*⌋, and *p_k_* = 0 for *k* > ⌊*σ*⌋ + 1, equaling *L*(*L* − 1)*A*^2^ − 2*C*(*L* − 1)*A* + *C*^2^ = (⌊*σ*⌋ + 1) ⌊*σ*⌋ − 2*σ*⌊*σ*⌋ + *σ*^2^.

If *σ* is an integer, then ⌊*σ*⌋ = ⌈*σ*⌉ = *σ*, and the maximum occurs with *p*_1_ = *p*_2_ = … = *p*_*σ*_ = 1, *p*_*σ*+1_ = *σ* − ⌊σ⌋ = 0, and *p_k_* = 0 for *k* > *σ* + 1, equaling *L*(*L* − 1)*A*^2^ − 2*C*(*L* − 1)*A* + *C*^2^ = ⌊*σ*⌋(⌊*σ*⌋ − 1) — 2*σ*(⌊*σ*⌋ − 1) + *σ*^2^.

In both cases, the maximum simplifies to (*σ* − ⌊σ⌋)^2^ + ⌊*σ*⌋, noting that ⌊*σ*⌋ = ⌊*σ*⌋ = *σ* in the latter case.

In our application of the theorem in the main text, we assume 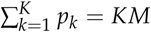, so that *KM* plays the role of *σ*. We thus obtain that the maximal value of 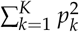 for sequences 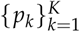 with *p_k_* ∈ [0,1], *k* < *ℓ* implies *p_k_* < *p_ℓ_*, and 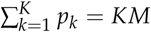 is (*KM* − ⌊*KM*⌋)^2^ + ⌊*KM*⌋, with equality if and only if *p_k_* = 1 for 1 ≤ *k* ≤ ⌊*Km*⌋, *p*_⌊*km*⌋+1_ = {*KM*}, and *p_k_* = 0 for *k* > ⌊*KM*⌋ + 1. Considering all sequences 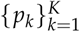 with *p_k_* ∈ [0,1] and not necessarily ordered such that *k* < ℓ implies *p_k_* < *p_ℓ_*, the maximum is achieved when any ⌊*KM*⌋ terms of the sequence equal 1, one term is {*KM*}, and all remaining terms are 0.

## Appendix B. Local minima of the upper bound on *F*

This appendix derives the positions and values of the local minima in the upper bound on *F* in terms of *M* (eq. 6).

### Positions of the local minima

To derive the positions of the local minima of the upper bound on *F* in terms of *M*, we study the function *Q_i_*(*x*) (eq. 7) on the interval [0,1) for *x*, where *i* = ⌊*KM*⌋ and *x* = *KM* − *i*, so that 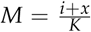. Recall that *K* and *i* are integers with *K* ≥ 2 and *i* in 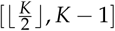. Note that *x* < 1 ensures that *M* < 1, in accord with our assumption of a polymorphic locus.

#### Theorem 2.

*Consider fixed integers K* ≥ 2 *and i in 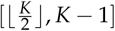*.

i. *Q_i_ (x) has no local minimum on [0,1) for K = 2 or for i = K − 1.*
ii. *For K ≥ 3 and i in 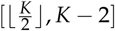, Q_i_(x) has a unique local minimum on the interval [0,1) for x, with position denoted x_min_.*
iii. *For odd K ≥ 3 and 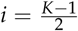, x_min_ = 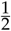.*
iv. *For all other (K, i) with K ≥ 3 and i in 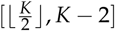, x_min_ = λ(K, i), where*

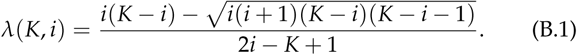

*Proof.* We take the derivative of *Q_i_*(*x*):

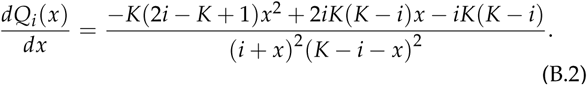

As 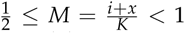, the denominator in eq. B.2 is non-zero, and *dQ_i_* (*x*)/*dx* = 0 is equivalent to a quadratic equation in *x*:

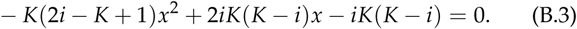

If 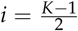, then the quadratic term in eq. B.3 vanishes and eq. B.3 becomes a linear equation in x, with solution *x* = 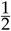. That the solution is a local minimum follows from the continuity of *Q_i_*(*x*) on [0,1) together with the fact that *Q_i_*(0) = *Q_i_*(1) = 1 and *Q_i_* (*x*) < 1 for 0 < *x* < 1. Consequently, if *K* is odd, then the local minimum for 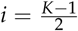 occurs at 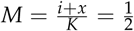, the lowest possible value of M. This establishes (iii).

Excluding 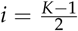 for odd *K*, for all 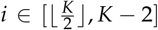 with *K* ≥ 3, eq. B.3 has a unique solution in [0,1); this solution has *x* = *λ*(*K,i*) (eq. B.1). The other root of eq. B.3 exceeds 1. That *x* = *λ*(*K,i*) represents a local minimum is again a consequence of the continuity of *Q_i_*(*x*) on [0,1) together with *Q_i_*(0) = *Q_i_*(1) = 1 and *Q_i_*(*x*) < 1 for 0 < *x* < 1. This establishes (ii) and (iv).

For the case of *i* = *K* − 1, eq. B.3 has a double root at *x* = 1, outside the permissible domain for *x*, [0,1). *Q_i_*(0) = 1, 0 ≤ *Q_i_*(*x*) ≤ 1 on [0,1), and *Q_i_*(*x*) approaches 0 as *x* → 1. Consequently, *Q_i_*(*x*) has no local minimum for *i* = *K* − 1. For *K* = 2, *i* = *K* − 1 is the only possible value of *i*, and *Q_i_*(*x*) has no local minimum. This establishes (i).

### Positions of the local minima for fixed *K* as a function of *i*

Having identified the locations of the local minima, we now explore how those locations change at fixed *K* with increasing *i*. For fixed *K* ≥ 3, we consider *x*_min_ from Theorem 2 as a function of *i* on the interval 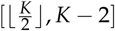. It is convenient to define interval *I*⁎;, equaling 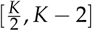 for even *K* and 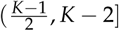 for odd *K*.

#### Proposition 1.

Consider a fixed integer K ≥ 3.

i. *The function x_min_(i) increases as i increases from 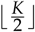 to K − 2.*
ii. *Its minimum is 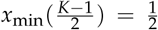 if K is odd, and 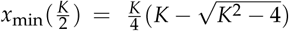 if K is even.*
iii. *Its maximum is x_min_ (1) = 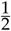 for K = 3, and for K > 3, it is*

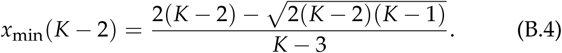

*Proof*. By Theorem 2, for fixed *K* ≥ 3 and *i* ∈ *I*⁎, *x*_min_(*i*) is given by eq. B.1. Treating *i* as continuous, we take the derivative:

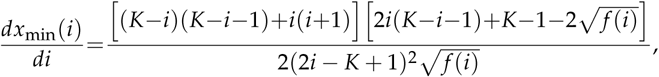

where *f* (*i*) = *i*(*i*+1)(*K*−*i*)(*K*−*i*−1). Because all other terms of *dx*_min_ (*i*)/*di* are positive for *i* in 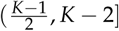, *dx*_min_(*i*)/*di* has the same sign as 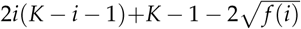.

Rearranging terms, we have

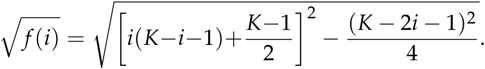

Because −(*K* − 2*i* − 1)^2^/4 < 0 for *i* in 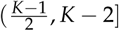,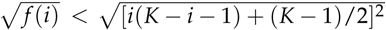, and 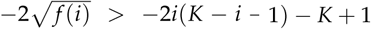. Consequently, 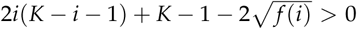 for all *i* in 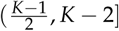. Thus, *dx*_min_(*i*)/*di* > 0 for all *i* in *I*⁎, and *x*_min_(*i*) increases with *i* in this interval.

For odd *K* and 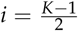, eq. B.1 gives lim_*i*→(*K* − 1)/2^+^_ λ(*K*, *i*) = 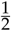. Thus, because 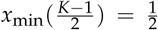 by Theorem 2, *x*_min_(*i*) is continuous at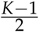. The function *x*_min_(*i*) therefore increases with *i* in the closed interval 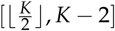. This proves (i).

Because *x*_min_(*i*) increases with *i* for all *i* in 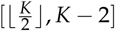, *x*_min_(*i*) is minimal when *i* is minimal. For *K* odd, the minimal value of *i* is 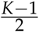. From Theorem 2, 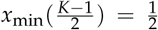 for all odd *K*. For even *K*, the minimal value of *i* is 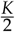. From Theorem 2, 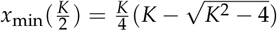. This proves (ii).

Similarly, *x*_min_(*i*) is maximal when *i* is maximal. From Theorem 2, the maximal value of *i* for which there exists a minimum of *Q_i_* (*x*) is *i*=*K*−2, and the position of this local minimum is 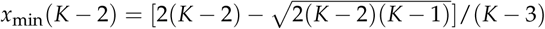. In particular, for 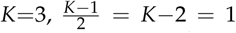, so there is a unique local minimum at position 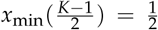. This proves (iii).

### Positions of the first and last local minima as functions of K

We now fix *i* and examine the effect of *K* on the local minimum at fixed *i*. We first focus on the interval closest to *M* = 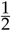—the *first local minimum* of the upper bound on *F*.

#### Proposition 2.

*Consider integers K* ≥ 3.

i. *For odd K, the relative position 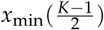 of the first local minimum does not depend on K and is 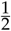.*
ii. *For even K, the relative position 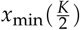 of the first local minimum decreases as K →∞, tends to 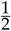, and is bounded above by 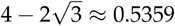.*

*Proof.* For odd *K*, the interval closest to 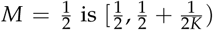. In this interval, from Proposition 1ii, the minimum occurs at 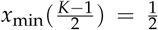 irrespective of *K*. This proves (i).

For even *K*, the interval for *M* closest to 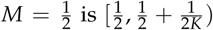. In this interval, from Proposition 1ii, the minimum has position 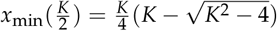. The derivative of this function is

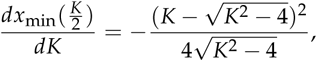

which is negative for all *K* ≥ 3. Thus, 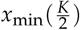 decreases with *K* for *K* ≥ 3. In addition, as 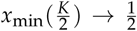. Because 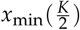 decreases with *K*, its maximum value is reached when *K* is minimal. The minimal even value of *K* is *K* = 4. Thus, 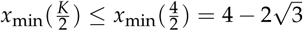. This proves (ii).

By Proposition 2, if *K* is large and even, then the first local minimum lies near the center of the interval 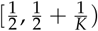 for *M*.

#### Proposition 3.

*For integers K ≥ 3, the relative position x_min_ (K − 2) of the last local minimum increases as K → ∞, and tends to 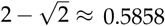.*

*Proof.* From Theorem 2, for *K* = 3 and *K* = 4, there is a single local minimum. Hence, from Proposition 2, the position of the last local minimum is *x*_min_ (1) = 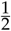 for *K* = 3 and 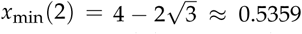 for *K* = 4. The position of the last local minimum then increases from *K* = 3 to *K* = 4.

If *K* > 3, from Proposition 1iii, the position of the last local minimum follows eq. B.4. We take the derivative

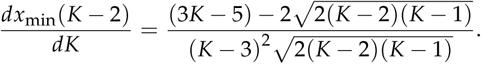

For *K* > 3, the denominator is positive, and *dx*_min_(*K* − 2)/*dK* has the same sign as its numerator. Because for *K* > 3, (3*K* − 5)^2^ − 8(*K* − 2)(*K* − 1) = (*K* − 3)^2^ > 0, we have 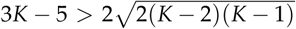 and a positive numerator. Then *dx*_min_ (*K* − 2) /*dK* > 0, and *x*_min_ (*K* − 2) increases for *K* > 3.

From eq. B.4, *x*_min_ (*K* −2) tends to 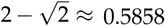 as *K* → ∞. Thus, the last local minimum is not at the center of interval *I*_*K*−2;_ rather, it is nearer to the upper endpoint. Because *x*_min_ (*K* − 2) increases with *K*, *x*_min_(*K* − 2) < lim_*K*→∞_ *x*_min_(*K* − 2), and the last local maximum has position bounded above by 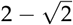.

As we have shown in Proposition 1i that for fixed *K*, as *i* increases from 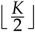 to *K*−2, the relative position of the local minimum increases, this relative position is restricted in the interval 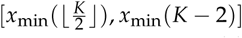. Further, because from Proposition 2, 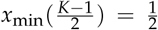 for odd *K* and 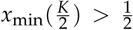 for even *K*, and from Proposition 3, 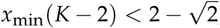, the relative positions of the local minima must be in the interval [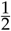,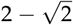).

Figure B1 illustrates as functions of *K* the relative positions of the first local minimum 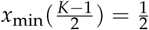 for *K* odd, 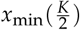 for *K* even) and the last local minimum (*x*_min_(*K* − 2)). The restriction of these positions to the interval [ 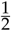,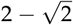) is visible, with the first local minimum lying closer to the center of interval [0,1) for *x* than the last local minimum. The decrease in the position of the first local minimum for even *K* alternating with values of 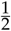 for odd *K* (Proposition 2) and the increase in the position of the last local minimum (Proposition 3) are visible as well.

**Figure B1.**
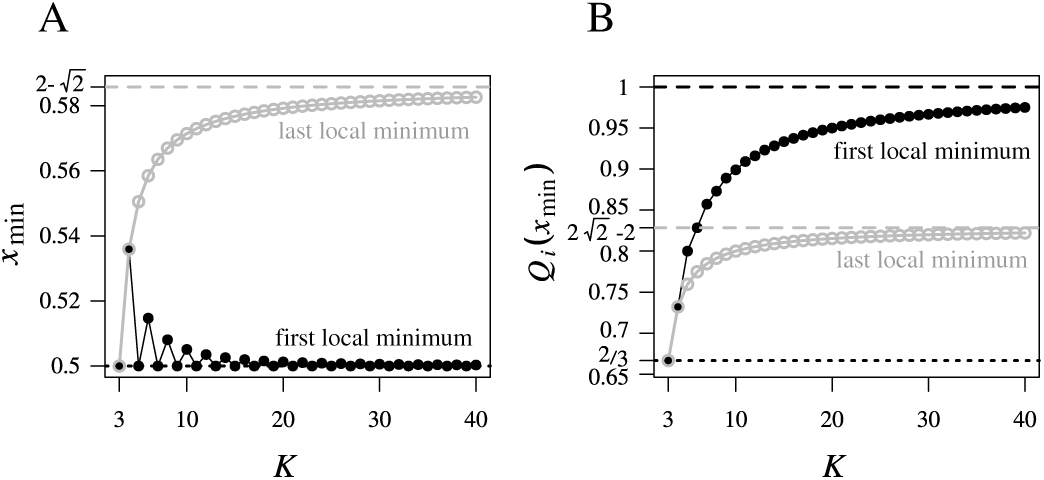
The first and last local minima of *F* as functions of the frequency *M* of the most frequent allele, for *K* ≥ 3 sub-populations. (A) Relative positions within the interval 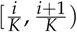 of the first and last local minima, as functions of *K*. The position *x*_min_ (*i*) of the local minimum in interval *I_i_* is computed from eq. B.1. If *K* is odd, then this position is 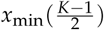; if *K* is even, then it is 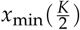. The position of the last local minimum is *x*_min_(*K* − 2). Dashed lines indicate the smallest value for *x*_min_ (*i*) of 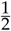, and the limiting largest value of 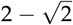. (B) The value of the upper bound on *F* at the first and last local minima, as functions of *K*. These values are computed from eq. 6, taking ⌊*KM*⌋ = *i* and {*KM*} = *x*_min_(*i*), with *x*_min_(*i*) as in part (A). Dashed lines indicate the limiting values of 1 and 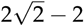 for the first and last local minima, respectively.

### Values at the local minima

We obtain the value of the local minima of the upper bound on *F* in each interval *I_i_* by substituting into eq. 7 the value of *i* for interval *I_i_* and its associated *x*_min_(*i*) from Theorem 2. We obtain

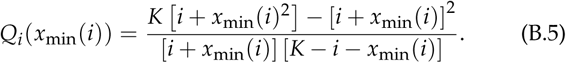

Note that for odd *K*, although λ(*K*, *i*) is undefined at 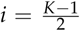, *x*_min_(*i*) is continuous. Thus, *Q_i_*(*x*_min_(*i*)) is also defined and continuous for all 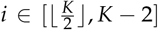. We consider *Q_i_* as a function of *i* on this interval.

#### Proposition 4.

For fixed *K* ≥ 3, the local minima *Q_i_*(*x*_min_(*i*)) decrease as *i* increases from 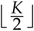 to *K* − 2.

Proof. We take the derivative *dQ_i_*(*x*_min_ (*i*))/*di* for fixed *K* and *i* ∈ *I*⁎. From eqs. B.5 and B.1,

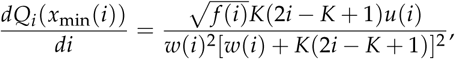

where 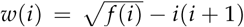 and 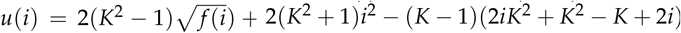.

For all *i* ∈ *I*⁎, the denominator of the derivative is positive, as are 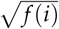, *K*, and 2*i* − *K* + 1. Hence, *dQ_i_*(*x*_min_(i))/*di* has the same sign as *u* (*i*).

Because *f*(*i*) decreases for 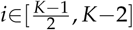, 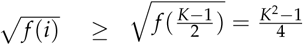, and 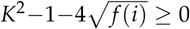. We factor *u*(*i*):

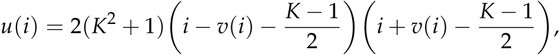

where 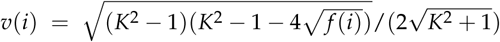. Because *v*(*i*) ≥ 0, for all 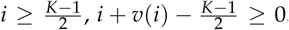. Thus, the sign of *u*(*i*) is given by the sign of

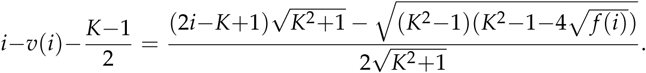

for 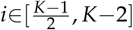

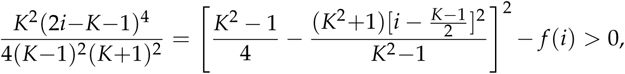

and hence,

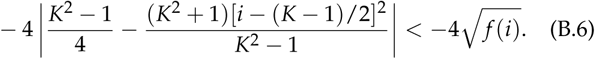

The term [*i* − (*K* − 1)/2]^2^ increases as a function of *i* for 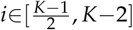. Hence, 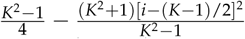 decreases with *i*. It is minimal at the largest value in the permissible domain for *i*, or *i* = *K* − 2, with minimum 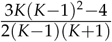. The denominator of this quantity is positive and the numerator increases with *K*. It is thus minimal for *K* = 3, at which 3*K*(*K* − 1)^2^ − 4 = 32 > 0. This proves that 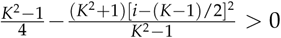 for 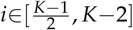.

We can then remove the absolute value in eq. B.6, and rearrange to obtain 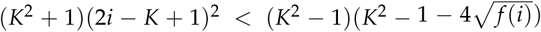. Both sides of this inequality are positive for 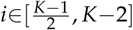, and we can take the square root to obtain 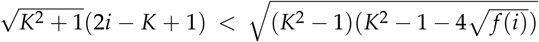. Hence 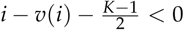 for 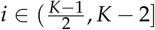, *dQ_i_*(*x*_min_(*i*))/*di* < 0 for *i* ∈ *I*⁎, and the local minima *Q_i_*(*x*_min_(*i*)) decrease with *i*.

#### Proposition 5.

*For K = 3, the first local minimum Q_i_(x_min_(i)) has value 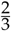; for K ≥ 3, the first local minimum increases as a function of K and tends to 1 as K → ∞.*

*Proof.* From Proposition 2*i*, for *K* odd, the first local minimum is reached for 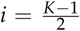 and *x* = 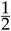, and the upper bound on *F* is 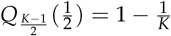. Thus, for *K* = 3, the first local minimum has value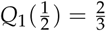. For even *K*, the first local minimum is reached if 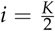 and 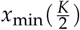, with upper bound on *F* equal to

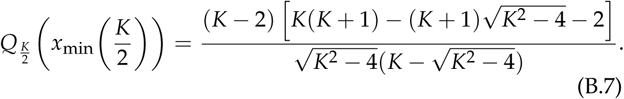

Denote by 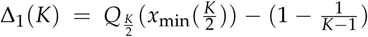 the difference between the first local minimum for even *K* and the first local minimum for odd *K* − 1, and by 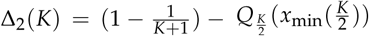 the difference between the first local minimum for odd *K* + 1 and the first local minimum for even *K*. To show that the first local minimum increases with *K*, we must show that for all even *K* ≥ 4, (i) Δ_1_(*K*) > 0, and (ii) Δ_2_(*K*) > 0.

For (i), subtracting 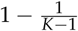 from eq. B.7, we have

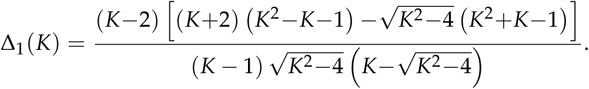

Because all other terms are positive for *K* ≥ 3, Δ_1_(*K*) has the same sign as 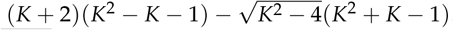. Dividing by 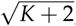, this quantity in turn has the same sign as 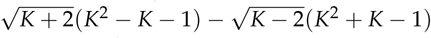. This last quantity is positive for *K* ≥ 4, as when we multiply it by the positive 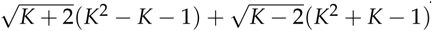, the result reduces simply to the number 4. This proves (i).

For (ii), subtracting eq. B.7 from 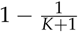, we have

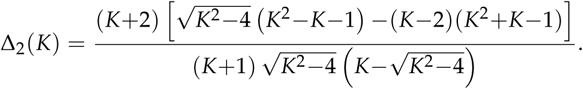

Because all other terms are positive for *K* ≥ 3, Δ_2_ (*K*) has the same sign as 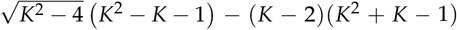. Dividing by 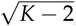, this quantity has the same sign as 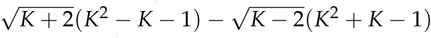, which was shown to be positive in the proof of (i). This demonstrates (ii).

From (i) and (ii), the value of the upper bound on *F* at the first local minimum increases with *K* for all *K* ≥ 3. To see that the limiting value is 1 as *K* → ∞, we note that the subsequence of values 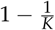 at odd *K* tends to 1 as *K* → ∞. As the sequence of values of the first local minimum with increasing *K* is monotonic and bounded above by 1, it is therefore convergent; as it has a subsequence converging to 1, the sequence converges to 1.

#### Proposition 6.

*For K = 3, the last local minimum Q_i_(x_min_(i)) has value 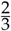; for K ≥ 3, the last local minimum increases as a function of K and tends to 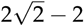 as K → ∞.*

*Proof.* From Proposition 5, for *K* = 3, the single local minimum has value 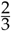. By Theorem 2, for *K* > 3, the last local minimum is reached when *i* = *K* − 2 and *x* = *x*_min_ (*K* − 2), in which case from eqs. B.5 and B.1, the upper bound on *F* is

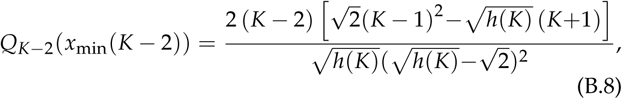

where *h*(*K*) = (*K* − 2)(*K* − 1).

We examine the derivative of *Q*_*K*−2_(*x*_min_(*K* − 2)) with respect to *K*. For *K* ≥ 3, *h*(*K*) > 0, and

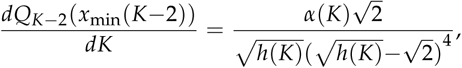

where 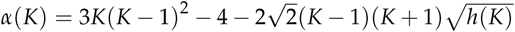. For *K* > 3, the derivative has the same sign as *α* (*K*).

We know that,

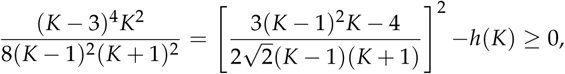

with equality only at *K* = 3. Because 3*K*(*K* − 1)^2^ − 4 is positive for 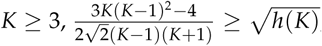 with equality only at *K* =3. Thus, *α*(*K*) > 0 and *dQ*_*K*−2_(*x*_min_(*K* − 2))/*dK* > 0 for *K* > 3, and hence, the last local minimum increases with *K*.

For the limit as *K* → ∞, we take the limit of *Q*_*K*−2_(*x*_min_(*K* − 2)) in eq. B.8, obtaining 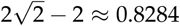.

## Appendix C. Computing the mean range of F

This appendix provides the computation of the integral *A*(*K*) (eq. 9) and its asymptotic approximation *Ã*(*K*).

### Computing A(K)

From eq. 9, we can compute *A*(*K*) by a sum over intervals. We use a partial fraction decomposition. For 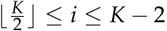,

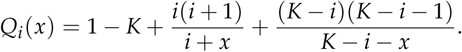

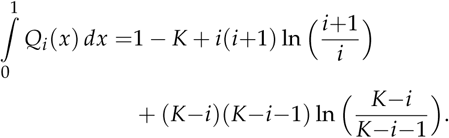

For *i* =*K* − 1,

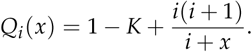

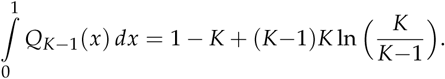

For 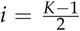

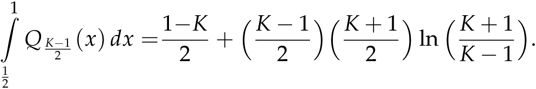

Thus, when *K* is even,

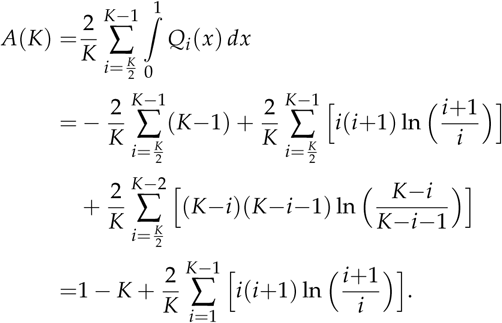

When *K* is odd,

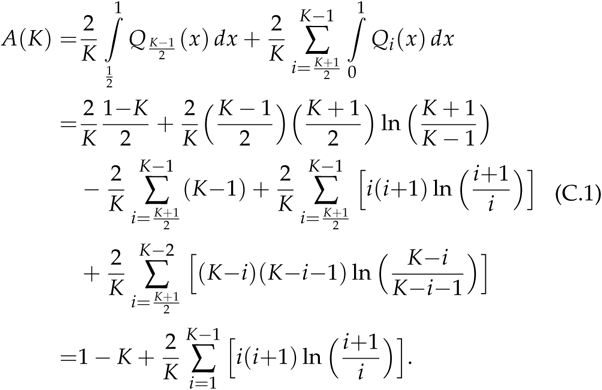

The expressions for *A*(*K*) are equal for even and odd *K*. We can simplify further:

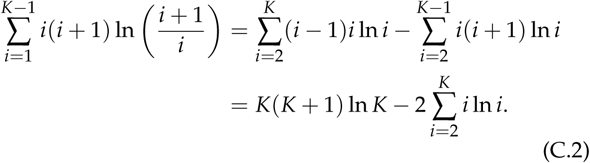

Substituting the expression from eq. C.2 into eq. C.1 and simplifying, we obtain eq. 10.

### Asymptotic approximation for A(K) (eqs. 11 and 12)

To asymptotically approximate *A*(*K*), we first need a large-*K* approximation of 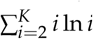. Because 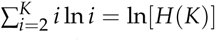, where 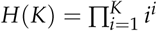 is the hyperfactorial function, we can use classical results about the asymptotic behavior of *H*(*K*):

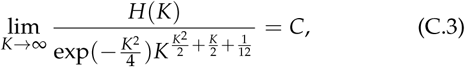

where C is the Glaisher-Kinkelin constant. Because the logarithm function is continuous at *C*, 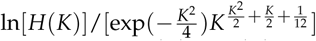 has limit ln *C* as *K* → ∞. Thus, if we write *f*(*K*) ~ *g*(*K*) for two functions that satisfy lim_*K→∞*_[*f* (*K*)/*g*(*K*)] = 1, then

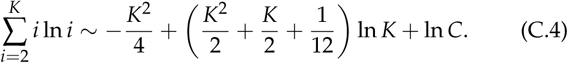

Substituting the expression from eq. C.4 into eq. 10, we obtain eq. 11 and function *Ã*(*K*) (eq. 12).

### Monotonicity of eq. 10 in K

#### Theorem 3.

*A(K) increases monotonically in K for K ≥ 2.*

*Proof*. We must show that Δ*A*(*K*) = *A*(*K* + 1) − *A*(*K*) > 0 for all *K* ≥ 2. From the expression for *A*(*K*) in eq. 10, we have:

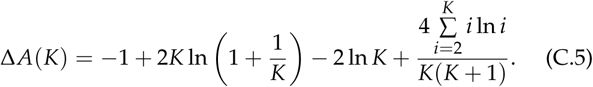

To show that Δ*A*(*K*) > 0, we find a lower bound for Δ*A*(*K*), denoted *D*(*K*), and then show that *D*(*K*) > 0.

We first find a lower bound for 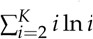. From the Euler-Maclaurin summation formula, we have

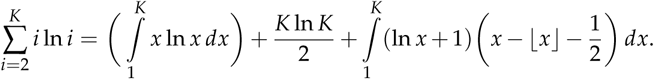

For all positive integers *i* ≥ 2,

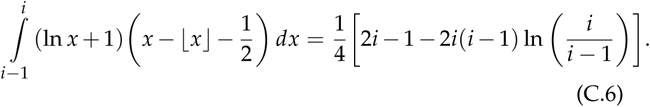

This integral can be seen to be positive by employing the equivalence for *i* > 1 of 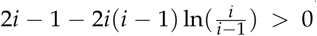 with 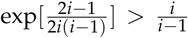. This latter inequality follows from the inequality 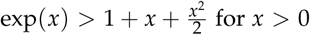 from the Taylor expansion of *e^x^*, noting that 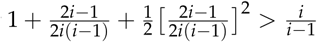.

Consequently, as the integral in eq. C.6 is positive for each 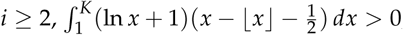, and

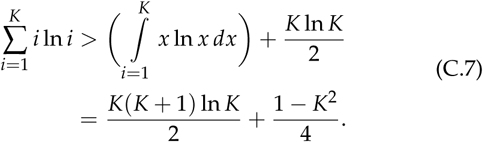

As a result, the following function is a lower bound for Δ*A*:

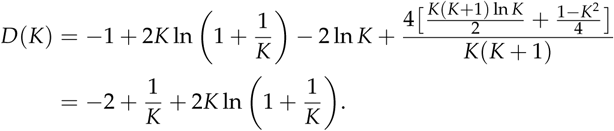

Dividing by 2*K* and substituting 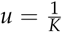, for *K* > 0, *D*(*K*) > 0 if and only if 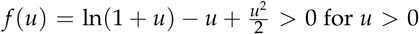for *u* >0. It can be seen that this latter inequality holds by noting that *f*(0) = 0 and 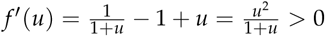

## Literature Cited

1000 Genomes Project Consortium et al., 2012 An integrated map of genetic variation from 1,092 human genomes. Nature 491: 56–65.

Akey, J. M., G. Zhang, K. Zhang, L. Jin, and M. D. Shriver, 2002 Interrogating a high-density SNP map for signatures of natural selection. Genome Res. 12: 1805–1814.

Alcala, N., J. Goudet, and S. Vuilleumier, 2014 On the transition of genetic differentiation from isolation to panmixia: what we can learn from G_ST_ and D. Theor. Pop. Biol. 93: 75–84.

Algee-Hewitt, B. F. B., M. D. Edge, J. Kim, J. Z. Li, and N. A. Rosenberg, 2016 Individual identifiability predicts population identifiability in forensic microsatellite markers. Curr. Biol. 26: 935–942.

Balloux, F., H. Brünner, N. Lugon-Moulin, J. Hausser, and J. Goudet, 2000 Microsatellites can be misleading: an empirical and simulation study. Evolution 54: 1414–1422.

Beaumont, M. A. and R. A. Nichols, 1996 Evaluating loci for use in the genetic analysis of population structure. Proc. R. Soc. Lond. Ser. B Biol. Sci. 263: 1619–1626.

Bonhomme, M., C. Chevalet, B. Servin, S. Boitard, J. Abdallah, S. Blott, and M. SanCristobal, 2010 Detecting selection in population trees: the Lewontin and Krakauer test extended. Genetics 186: 241–262.

Cann, H. M., C. De Toma, L. Cazes, M.-F. Legrand, V. Morel, L. Piouffre, J. Bodmer, W. F. Bodmer, B. Bonne-Tamir, A. Cambon-Thomsen, et al., 2002 A human genome diversity cell line panel. Science 296: 261–262.

Cañon, J., M. L. Checa, C. Carleos, J. L. Vega-Pla, M. Vallejo, and S. Dunner, 2000 The genetic structure of Spanish Celtic horse breeds inferred from microsatellite data. Anim. Genet. 31: 39–48.

Cornuet, J.-M., F. Santos, M. A. Beaumont, C. P. Robert, J.-M. Marin, D. J. Balding, T. Guillemaud, and A. Estoup, 2008 Inferring population history with DIY ABC: a user-friendly approach to approximate Bayesian computation. Bioinformatics 24: 2713–2719.

Edge, M. D. and N. A. Rosenberg, 2014 Upper bounds on *F_ST_* in terms of the frequency of the most frequent allele and total homozygosity: the case of a specified number of alleles. Theor. Pop. Biol. 97: 20–34.

Foll, M. and O. Gaggiotti, 2008 A genome-scan method to identify selected loci appropriate for both dominant and codominant markers: a Bayesian perspective. Genetics 180: 977–993.

Frankham, R., J. D. Ballou, and D. A. Briscoe, 2002 Introduction to Conservation Genetics. Cambridge University Press, Cambridge.

Günther, T. and G. Coop, 2013 Robust identification of local adaptation from allele frequencies. Genetics 195: 205–220.

Hartl, D. L. and A. G. Clark, 1997 Principles of Population Genetics. Sinauer, Sunderland, MA.

Hedrick, P. W., 1999 Highly variable loci and their interpretation in evolution and conservation. Evolution 53: 313–318.

Hedrick, P. W., 2005 A standardized genetic differentiation measure. Evolution 59: 1633–1638.

Hinds, D. A., L. L. Stuve, G. B. Nilsen, E. Halperin, E. Eskin, D. G. Ballinger, K. A. Frazer, and D. R. Cox, 2005 Whole-genome patterns of common DNA variation in three human populations. Science 307: 1072–1079.

Holsinger, K. E. and B. S. Weir, 2009 Genetics in geographically structured populations: defining, estimating and interpreting *F_ST_*. Nature Rev. Genet. 10: 639–650.

Hudson, R. R., 2002 Generating samples under a Wright–Fisher neutral model of genetic variation. Bioinformatics 18: 337–338.

International HapMap Consortium, 2005 A haplotype map of the human genome. Nature 437: 1299–1320.

Jakobsson, M., M. D. Edge, and N. A. Rosenberg, 2013 The relationship between *F_ST_* and the frequency of the most frequent allele. Genetics 193: 515–528.

Jost, L., 2008 G_ST_ and its relatives do not measure differentiation. Mol. Ecol. 17: 4015–4026.

Kim, T. H., K. S. Kim, B. H. Choi, D. H. Yoon, G. W. Jang, K. T. Lee, H. Y. Chung, H. Y. Lee, H. S. Park, and J. W. Lee, 2005 Genetic structure of pig breeds from Korea and China using microsatellite loci analysis. J. Anim. Sci. 83: 2255–2263.

Lawson Handley, L. J., K. Byrne, F. Santucci, S. Townsend, M. Taylor, M. W. Bruford, and G. Hewitt, 2007 Genetic structure of European sheep breeds. Heredity 99: 620–631.

Leinonen, T., R. J. S. McCairns, R. B. O’Hara, and J. Merilä, 2013 *Q_ST_*–*F_ST_* comparisons: evolutionary and ecological insights from genomic heterogeneity. Nature Rev. Genet. 14: 179–190.

Lewontin, R. and J. Krakauer, 1973 Distribution of gene frequency as a test of the theory of the selective neutrality of polymorphisms. Genetics 74: 175–195.

Li, J. Z., D. M. Absher, H. Tang, A. M. Southwick, A. M. Casto, S. Ramachandran, H. M. Cann, G. S. Barsh, M. Feldman, L. L. Cavalli-Sforza, et al., 2008 Worldwide human relationships inferred from genome-wide patterns of variation. Science 319: 1100–1104.

Long, J. C. and R. A. Kittles, 2003 Human genetic diversity and the nonexistence of biological races. Hum. Biol. 75: 449–471.

Maruki, T., S. Kumar, and Y. Kim, 2012 Purifying selection modulates the estimates of population differentiation and confounds genome-wide comparisons across single-nucleotide polymorphisms. Mol. Biol. Evol. 29: 3617–3623.

Maruyama, T., 1970 Effective number of alleles in a subdivided population. Theor. Pop. Biol. 1: 273–306.

Nei, M., 1973 Analysis of gene diversity in subdivided populations. Proc. Natl. Acad. Sci. USA 70: 3321–3323.

Pemberton, T. J., D. Absher, M. W. Feldman, R. M. Myers, N. A. Rosenberg, and J. Z. Li, 2012 Genomic patterns of homozygosity in worldwide human populations. Am. J. Hum. Genet. 91: 275–292.

Reddy, S. B. and N. A. Rosenberg, 2012 Refining the relationship between homozygosity and the frequency of the most frequent allele. J. Math. Biol. 64: 87–108.

Rosenberg, N. A. and M. Jakobsson, 2008 The relationship between homozygosity and the frequency of the most frequent allele. Genetics 179: 2027–2036.

Rosenberg, N. A., L. M. Li, R. Ward, and J. K. Pritchard, 2003 Informativeness of genetic markers for inference of ancestry. Am. J. Hum. Genet. 73: 1402–1422.

Rosenberg, N. A., S. Mahajan, C. Gonzalez-Quevedo, M. G. B. Blum, L. Nino-Rosales, V. Ninis, P. Das, M. Hegde, L. Molinari, G. Zapata, et al., 2006 Low levels of genetic divergence across geographically and linguistically diverse populations from India. PLoS Genet. 2: e215.

Rosenberg, N. A., J. K. Pritchard, J. L. Weber, H. M. Cann, K. K. Kidd, L. A. Zhivotovsky, and M. W. Feldman, 2002 Genetic structure of human populations. Science 298: 2381–2385.

Slatkin, M., 1985 Rare alleles as indicators of gene flow. Evolution 39: 53–65.

Tishkoff, S. A., F. A. Reed, F. R. Friedlaender, C. Ehret, A. Ranciaro, A. Froment, J. B. Hirbo, A. A. Awomoyi, J.-M. Bodo, O. Doumbo, M. Ibrahim, A. T. Juma, M. J. Kotze, G. Lema, J. H. Moore, H. Mortensen, T. B. Nyambo, S. A. Omar, K. Powell, G. S. Pretorius, M. W. Smith, M. A. Thera, C. Wambebe, J. L. Weber, and S. M. Williams, 2009 The genetic structure and history of Africans and African Americans. Science 324: 1035–1044.

Wakeley, J., 1998 Segregating sites in Wright’s island model. Theor. Pop. Biol. 53: 166–174.

Wakeley, J., 1999 Nonequilibrium migration in human history. Genetics 153: 1863–1871.

Wang, J., 2015 Does G_ST_ underestimate genetic differentiation from marker data? Mol. Ecol. 24: 3546–3558.

Weir, B. S., 1996 Genetic Data Analysis II. Sinauer, Sunderland, MA.

Whitlock, M. C., 2011 *G′_ST_* and D do not replace *F_ST_*. Mol. Ecol. 20: 1083–1091.

Wilkinson-Herbots, H. M., 1998 Genealogy and subpopulation differentiation under various models of population structure. J. Math. Biol. 37: 535–585.

Wright, S., 1951 The genetical structure of populations. Ann. Eugen. 15: 323–354.

